# Physiologically-based pharmacokinetic model for CAR-T cells delivery and efficacy in solid tumors

**DOI:** 10.1101/2025.08.20.671229

**Authors:** Andreas G. Hadjigeorgiou, Lance L. Munn, Triantafyllos Stylianopoulos, Rakesh K. Jain

**Affiliations:** Cancer Biophysics Laboratory, Department of Mechanical and Manufacturing Engineering, University of Cyprus; Edwin L. Steele Laboratories, Department of Radiation Oncology, Massachusetts General Hospital and Harvard Medical School, Boston, MA, USA

**Keywords:** vascular normalization, stroma normalization, CAR-T cells, mathematical model

## Abstract

Abnormal blood vessels limit the delivery and function of endogenous T cells as well as adoptively transferred Chimeric Antigen Receptor (CAR)-T cells in the tumor microenvironment (TME). We recently showed that vascular normalization using anti-VEGF therapy can overcome these challenges and improve the outcome of CAR-T therapy in glioblastoma models in mice. Here, we developed a physiologically based pharmacokinetic model to simulate the dynamics of both adoptively transferred CAR-T cells and endogenous immune cells in solid tumors following vascular normalization. Similar to our data, our model simulations show that vascular normalization reprograms the TME from immunosuppressive to immunosupportive—enhancing infiltration of endogenous CD8⁺ T cells and CAR-T cells, increasing M1 macrophages, and reducing M2 macrophages and regulatory T cells—thereby improving efficacy. Strikingly, vascular normalization reduces the number of infused CAR-T cells needed for tumor control by an order of magnitude. Moreover, synchronizing a second CAR-T infusion at their peak proliferative phase maximizes antitumor function. Furthermore, the efficacy of CAR-T cells engineered to secrete anti-VEGF antibody depends on the ability of CAR-T cells to induce vascular normalization. Additionally, combining vascular and stromal normalization can improve the efficacy of anti-VEGF antibody-producing FAP-CAR-T cells for the treatment of desmoplastic tumors such as pancreatic ductal adenocarcinoma. Finally, the model predicts that local CAR-T delivery can sustain high concentrations within the TME and induce recruitment of other antitumor immune cells, improving outcomes. Our model provides a versatile framework to optimize dosing strategies, treatment sequencing, and delivery routes for improving CAR-T therapies for solid tumors.

**Significance Statement**

Preclinical studies and early clinical trials of CAR-T therapy show encouraging responses in glioblastoma, diffuse midline gliomas, and neuroblastoma, yet substantial obstacles remain for effective CAR-T therapy for solid tumors. Building on our discovery that judicious VEGF blockade normalizes tumor vessels and enhances CD8⁺T-cell infiltration, we developed a mathematical model to optimize CAR-T therapy for solid tumors. Simulations predict that vascular normalization can render the TME immunosupportive and decrease CAR-T doses tenfold. In desmoplastic tumors, FAP-CAR-T efficacy is improved by combining anti-VEGF and stromal normalizing agents. Optimal scheduling and direct intratumoral delivery can mitigate T-cell exhaustion and improve tumor control further. Thus, our model serves as a strategic roadmap for optimal CAR-T deployment in solid tumors.

## Introduction

Glioblastoma (GBM) is a uniformly fatal malignancy with limited treatment options and a median survival of less than two years (1). While Chimeric Antigen Receptor (CAR)-T cell therapy has revolutionized the treatment of hematological malignancies (2–4), its efficacy against solid tumors, such as GBM, remains dismal (5–9) (see Table 1 for the indicative clinical trials and their key results). Our hypothesis is that for CAR-T cell therapy to be efficacious in solid tumors, systemically or locally injected CAR-T cells need to accumulate in the tumor microenvironment (TME) in sufficient quantities and remain activated to induce tumor regression without causing systemic toxicity. However, the abnormal TME not only hinders the uniform delivery of CAR-T but also impairs their function even after these cells accrue in the TME.

**Table 1.**
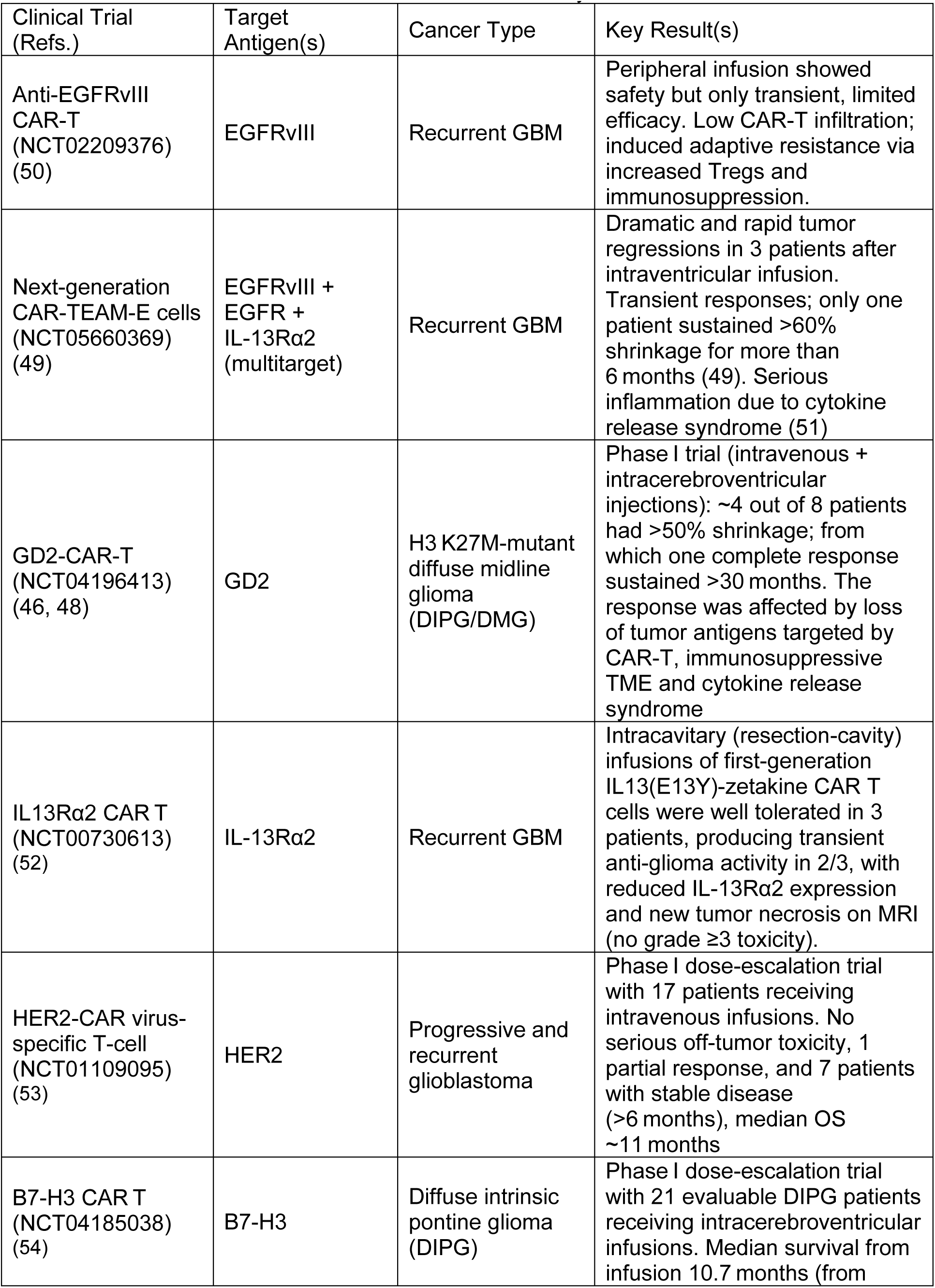
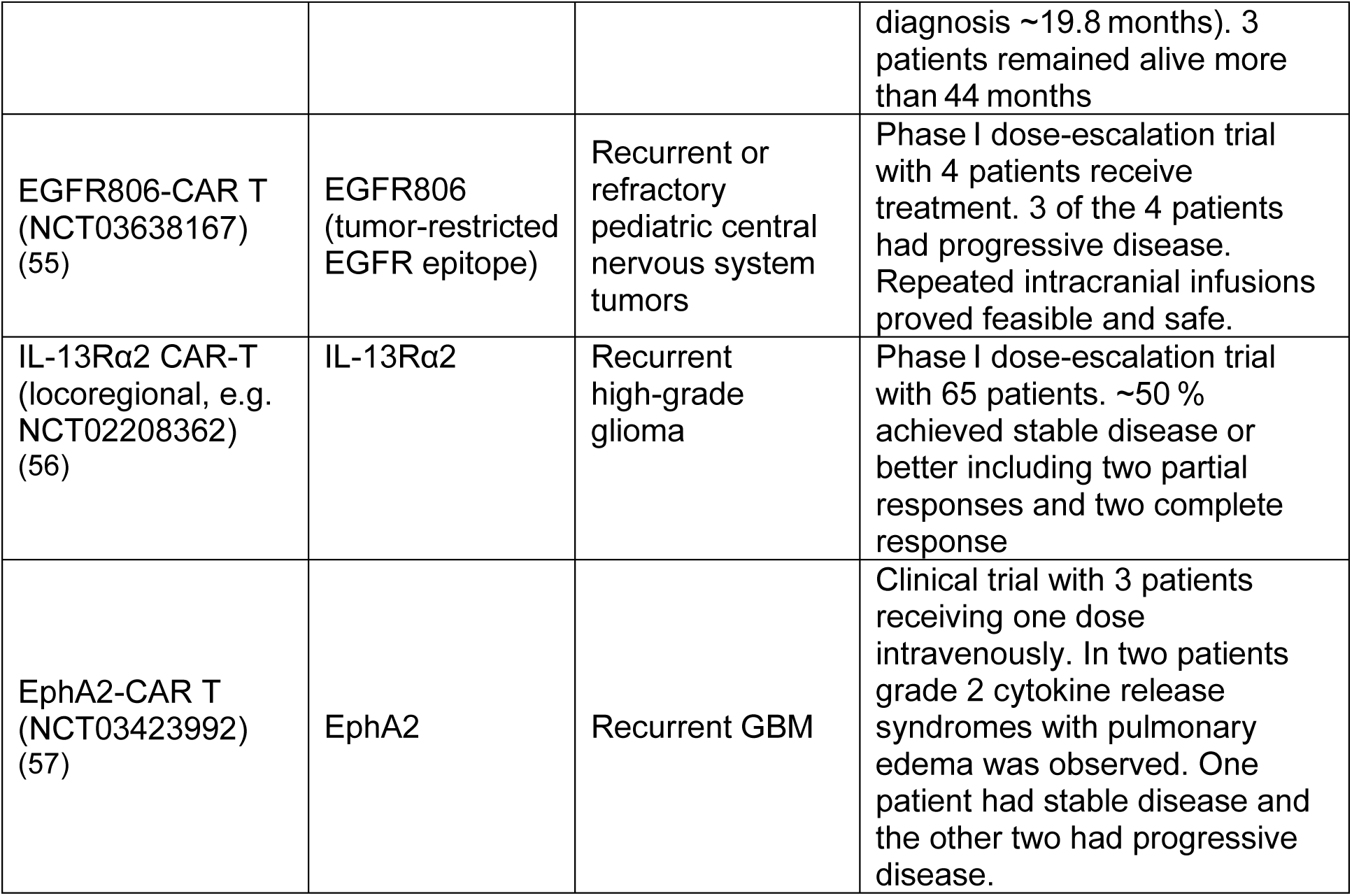
Clinical trials of CAR-T cells in solid tumors and their key results.

The TME encompasses blood and lymphatic vessels, stromal cells such as fibroblasts, and other non-neoplastic components such as immune cells (both pro- and anti-tumor)– all embedded in an extracellular matrix (ECM) (10). Unlike normal vessels, the tumor vessels are abnormal in structure and function (e.g., hyperpermeable) due to an imbalance between pro-angiogenic factors, such as VEGF and anti-angiogenic factors, such as thrombospondin (11, 12) (Fig. 1, bottom left panel).

**Figure 1.**
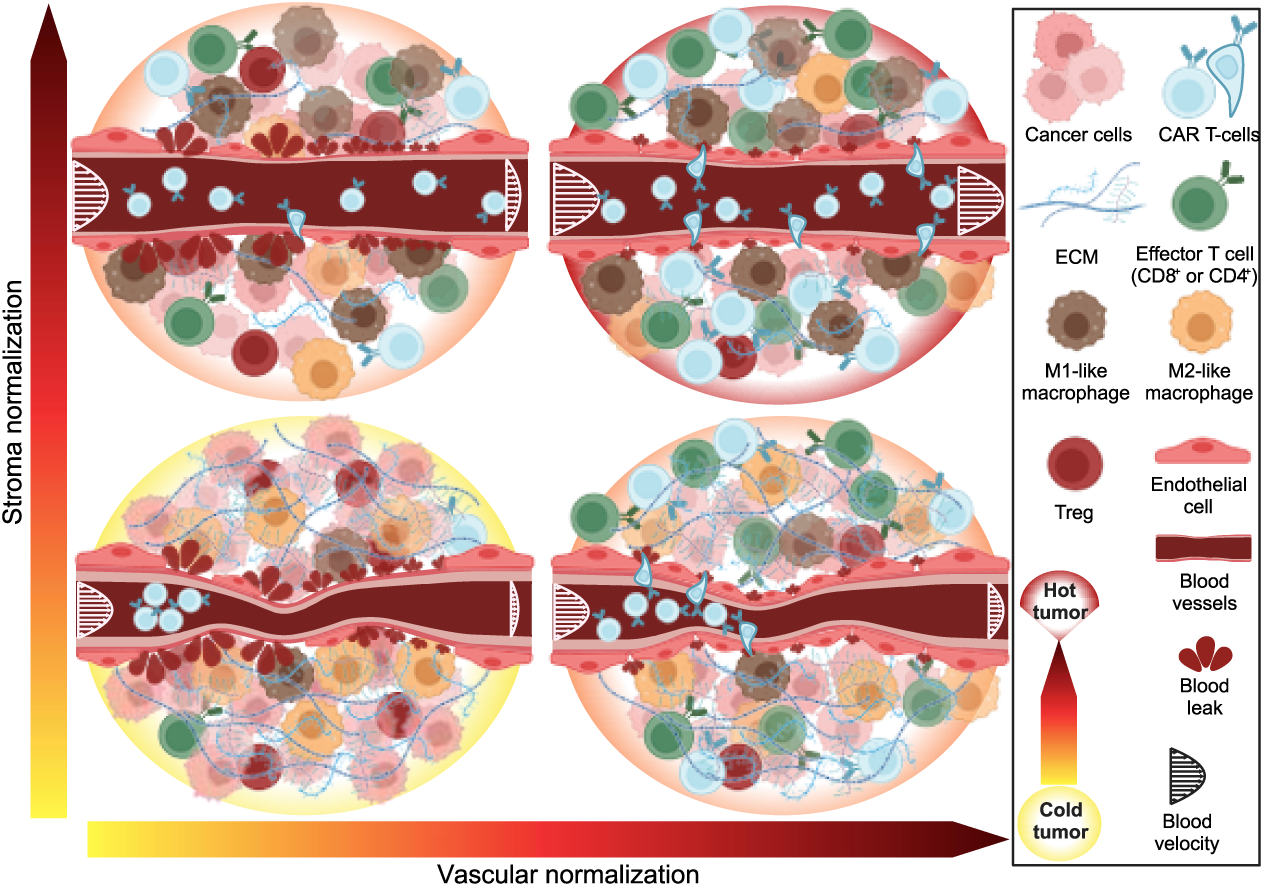
Vascular and stroma normalization in solid tumors. The TME of a cold tumor (bottom left) is characterized by excessive ECM and compressed and hyperpermeable vessels. These abnormal vessels cause immunosuppression with abundant immunosuppressive cells such as M2-like macrophages and Treg cells and few immunosupportive M1-like macrophages, effector T cells and CAR-T cells. Stroma normalization can normalize ECM levels, decompressing the vessels and partially restoring perfusion. Vascular normalization with anti-VEGF treatment (bottom right) fortifies the blood vessels wall, reducing permeability (leakiness) and thus, increasing their perfusion. The combination of these two normalization strategies (upper right) can optimally restore perfusion in solid tumors allowing the transport and infiltration of CAR-T cells into the tumor to create a hot TME with sufficient immunosupportive immune cells.

Furthermore, in tumors, the overproduction of ECM components, mainly hyaluronan and collagen, increases intratumoral mechanical forces, causing compression of tumor blood vessels (13–15) (Fig. 1, bottom left panel). Vessel compression and leakiness lead to hypo-perfused and dysfunctional tumor vessels. These dysfunctional tumor vessels, combined with increased oxygen consumption and anaerobic glycolysis, contribute to the development of hypoxic and acidic TME. Hypoxia and low pH induce the production of immunosuppressive molecules, including TGF-β and VEGF (16, 17), and affect anti-tumor immune cell function in many ways (16, 17). These immunosuppressive cytokines also reduce the expression of adhesion molecules on the vascular endothelium, preventing endogenous immune cells and CAR-T cells from adhering to and migrating across the vessel walls into the tumor (17). Hypoxia also promotes the recruitment of immunosuppressive regulatory T cells and reprograms tumor-associated macrophages to a protumor phenotype (17).

To overcome these challenges, we proposed to “normalize” the tumor vessels using judicious doses of antiangiogenic agents (12, 18, 19). This strategy reduces the excessive leakiness and chaotic structure of the vessels, increases the adhesion molecules on the vessel wall, and reprograms the TME from immunosuppressive to immunostimulatory (12, 16, 17) (Fig. 1, bottom right panel). After revealing the molecular mechanism of vascular normalization and its benefits in a number of preclinical models, we verified its benefit in clinical trials for patients with GBM, breast, colorectal, liver, and lung cancer (20). Our preclinical finding that vascular normalization can improve the outcome of immunotherapy (21) has been confirmed by clinical trials combining anti-VEGF/R agents with immune-checkpoint blockers, and has formed the basis for 7 FDA approvals of such combinations for endometrial, kidney, lung, and liver cancers (16, 22). Of interest, many pharmaceutical companies are now developing bi-functional antibodies that target both PD1/L1 and VEGF/R, and one of these - Ivonescimab - has shown positive outcomes in a phase III clinical trial in lung cancer (23).

Vascular normalization can also improve the outcome of immunotherapies that involve the adoptive transfer of immune cells. In 2010, Rosenberg and colleagues demonstrated that anti-VEGF-induced vascular normalization improves the delivery and efficacy of adoptively transferred T cells in mice bearing B16 melanoma (24). Recently, we demonstrated the benefit of anti-VEGF induced vascular normalization in the outcome of EGFRviii-CAR-T cell therapy in two murine GBM models (25). Vascular normalization not only improves the delivery and function of EGFRviii-CAR-T cells but also increases the delivery and activation of endogenous T cells (25). Consequently, many academic and biotech companies are now developing CAR-T cells that can produce anti-VEGF scFv antibodies (26). Furthermore, stroma normalization can further increase vessel perfusion (27, 28) and has the potential to increase CAR-T cell infiltration by decompressing tumor vessels and increasing porosity within the tumor (Fig. 1, top left panel); stroma and vascular normalization (Fig. 1, top right panel) could also have synergistic effects. This could be beneficial in desmoplastic/fibrotic tumors, which have abundant collapsed vessels. All these developments raise important questions about the optimal design of these preclinical and clinical studies for both cranial and extracranial tumors.

To this end, we developed a physiologically based pharmacokinetics (PBPK) model for CAR-T cells as well as endogenous immune cells with particular emphasis on TME normalization. Our current model builds on first-generation PBPK models to describe the biodistribution of effector cells in normal and neoplastic mammalian tissues (29). As summarized in Table 2, early PBPK models (Zhu et al. (29), Melder et al. (30), Nikitich et al. (31)) simulated lymphocyte trafficking and biodistribution—revealing that pulmonary and splenic retention severely limits tumor delivery. Subsequent PK-PD and QSP frameworks expanded the scope to include tumor growth and treatment efficacy, antigen-binding dynamics, dosing-route variations, cytokine/CRS risk, and T-cell exhaustion. For example, Singh et al. (32) demonstrated that CAR-T expansion is driven more by tumor burden than dose; Jiang et al. (33) showed that local delivery near the tumor enhances infiltration and early proliferation; Li et al. (34) captured antigen-density–dependent killing kinetics; Adhikarla et al. (35) optimized sequencing for combined radionuclide therapy; Hardiansyah et al. (36) linked high tumor load to increased cytokines release syndrome risk; and Lai et al. (37) highlighted exhaustion dynamics as key determinants of tumor evolution.

**Table 2.** Summary of the relevant models and their findings.

| Relevant model (ref) | Model Components ✓/× | Main Findings |
| --- | --- | --- |
| Zhu et al. (1996) (29) | Trafficking/recirculation ✓, multiorgan biodistribution ✓, tumor compartment ✓, no explicit treatment efficacy ×, no tumor growth-response ×, no endogenous immune system × | Retention of effector cells in lungs and spleen significantly impedes therapeutic effectiveness. |
| Melder et al. (2002) (30) | Trafficking/recirculation ✓, multiorgan biodistribution ✓, tumor compartment ✓, no explicit treatment efficacy ×, no tumor growth-response ×, no endogenous immune system × | Lymphocyte retention in normal tissues limits initial tumor recruitment; splenectomy shifts organ distribution without improving tumor localization. |
| Hardiansyah et al. (2019) (36) | QSP model-PD of CAR-T therapy in hematologic malignancies ✓, cytokine response/exhaustion modeling ✓, tumor growth✓, explicit treatment efficacy✓, No multiorgan biodistribution ×, no endogenous immune system × | Higher tumor burden elicits a stronger cytokine response and increases the risk of severe cytokine release syndrome (CRS). |
| Singh et al. (2020) (32) | CAR T therapy in hematologic malignancies ✓, multiorgan biodistribution ✓, CAR affinity, expansion ✓, tumor growth✓, explicit treatment efficacy✓, no endogenous immune cells × | CAR T expansion in blood is more sensitive to tumor burden than to dose levels. |
| Jiang et al. (2020) (33) | Trafficking/recirculation ✓, multiorgan biodistribution ✓, hematologic malignancies ✓, tumor growth✓, explicit treatment efficacy✓, endogenous T cells ✓, no other endogenous immune cells ×, no CAR T therapy ×, | Local (near tumor) delivery enhances initial infiltration and earlier proliferation versus systemic routes. |
| Li et al. (2023) (34) | In vitro cell-level binding model ✓ multiple CAR-T to one tumor cell, tumor growth✓, explicit treatment efficacy✓ | In vitro model shows CAR T expansion rate and killing dynamics critically depend on antigen-density and multi binding. |
| Nikitich et al. (2024) (31) | Trafficking ✓, multiorgan biodistribution ✓, endogenous and exogenously administered T cells ✓, no tumor compartment ×, no explicit treatment efficacy ×, no tumor growth response × | Pronounced accumulation of T cells in the spleen, limiting delivery to tumor. |
| Adhikarla et al. (2024) (35) | CAR T therapy in multiple myeloma ✓, radionuclide therapy ✓ | Optimal sequencing and dosing strategies of CAR T |
|  | (sequencing/dose), tumor growth ✓, explicit treatment efficacy✓, no multiorgan biodistribution×, no endogenous immune cells × | therapy and radionuclide therapy. |
| Lai et al. (2025) (37) | Endogenous T cells ✓, tumor growth✓, T cells tumor cells interactions✓, exhaustion modeling ✓, no multiorgan biodistribution× | T cell exhaustion dynamics are critical determinants of tumor evolution and therapy outcome. |

The current model incorporates components of the endogenous immune system, immune activation, immune exhaustion and suppression, and the effects of TME normalization and CAR-T therapy. We use our model to track both CAR-T cells and key endogenous immune cell populations—dendritic cells, M1/M2 macrophages, regulatory T cells, naïve and effector CD8⁺ T cells, and APCs—across anatomically connected organs, blood, and lymphatic spaces (Fig. 2A). Each organ compartment is split into vascular and extravascular sub-compartments, with blood flow (red/blue arrows, Fig. 2A) and lymphatic drainage (green arrows, Fig. 2A) governing cell trafficking from one compartment to the other. The model considers the transport of all cells from the vascular to the extravascular space (Fig. 2B) and cell recirculation back to blood circulation through the lymphatic vessels (Fig. 2B). In the tumor compartment (Fig. 2C), tumor growth and immune–tumor interactions are explicitly modeled: CAR-T and effector CD8⁺ T cells cause cytolysis of tumor cells, while M1 macrophages and dendritic cells phagocytose tumor debris and mature into APCs (Fig. 2D), which then recirculate to lymph nodes to prime naïve CD8⁺ T cells. Conversely, M2 macrophages and Tregs suppress functions of APCs, CAR-T cells, and effector T cells (Fig. 2E). All transport and interaction kinetics are defined by population-balance equations detailed in the Supplementary Information, and the interactions between cells are provided in Supplementary Table 1.

**Figure 2.**
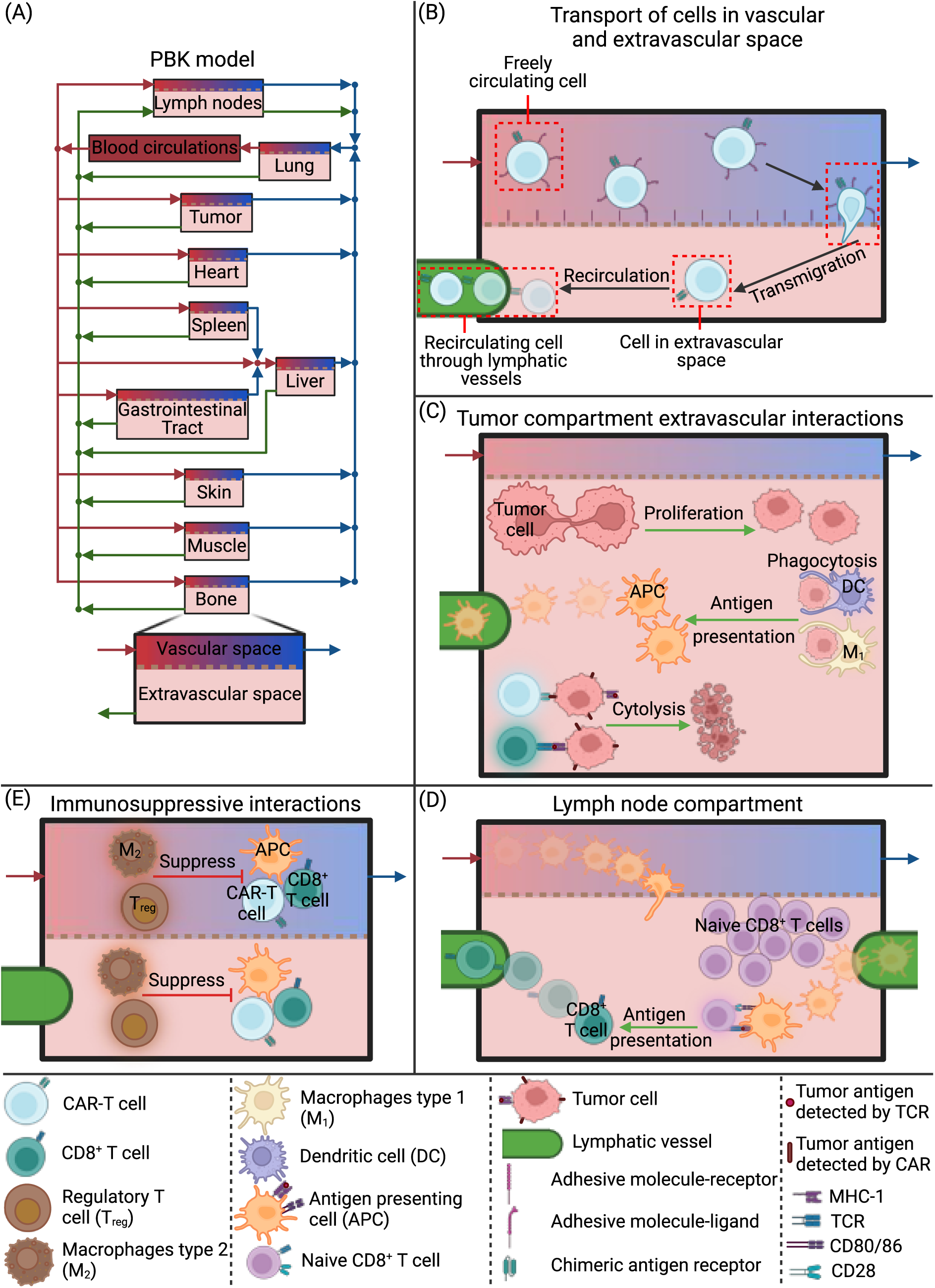
Schematic of the PBPK model used to simulate the immune response, tumor growth and CAR-T cell treatment with the main interactions in the compartments. A) The PBPK model with the compartments of the body which are connected by the blood circulation (blue and red arrows) and lymph circulation (green arrows). B) the transport of cells from the vascular to the extravascular space (transmigration) in the compartments and the recirculation of the cells through lymphatic vessels. C) The interactions in the extravascular space of the tumor compartment where the tumor cells proliferate, and other cells interact with them. D) Activation of naïve CD8^+^T cells in the lymph node compartment. E) The immunosuppressive interactions with regulatory T cells and macrophages type 2.

In this study, we first used our model to simulate combined anti-VEGF and CAR-T cell therapy by integrating our published in vivo experimental data on GBM (25, 38) as well as relevant data from the literature (24, 39). Then, we employ the model to evaluate multiple CAR-T treatment protocols under varying vascular conditions. These included systemic CAR-T injections at different doses and time points, both with and without prior anti-VEGF–induced vascular normalization. Dual systemic dosing strategies were also tested by varying the interval and timing of injections. Another scenario involved first inducing vascular normalization with anti-VEGF, followed by systemic delivery of anti-VEGF–secreting CAR-T cells, assessing the optimal timing and required CAR-T cell density to maintain normalization. We then simulated the effect of combining vascular and stromal normalization using anti-VEGF, agents targeting ECM, and FAP-CAR-T cells. Finally, we compared local versus systemic CAR-T delivery routes to determine their relative efficacy.

## Results

### Model calibration with experimental data

First, we calibrated the model by comparing its predictions with our published in vivo experimental data from mice bearing orthotopic syngeneic (GSC005) GBM murine tumors (25) and human (MGG8) GBM xenograft tumors (38). For the MGG8 tumors, we considered only the treatment groups that are relevant to our model: the control and anti-VEGF treatment groups (38). This experiment was performed with immunodeficient mice (38); thus, the CD8^+^ T cells were set to zero, as well as the related model parameters. The tumor growth data and infiltrated macrophages (determined by flow cytometry, percentages of M1 and M2 normalized to total macrophages) (38) were used to validate the model. The GSC005 experiment involved four treatment groups: control, anti-VEGF, CAR-T, and a combination of anti-VEGF and CAR-T cell (anti-VEGF + CAR-T) therapies (25). From this experiment, tumor growth data, infiltrated CAR-T cells measured by multiphoton microscopy (number of cells per volume), immunofluorescence staining (number of cells per volume) and flow cytometry (percentage of CAR-T cells normalize to total T cells) as well as measurements of CD8^+^ T cells and FOXP3^+^ CD4^+^ T cells (Tregs) by flow cytometry (percentage of T cells over total T cells) (25) were employed for validation.

With this procedure, we achieved general agreement between the model and experimental data (**Fig. 3**). Fig. 3A demonstrates the comparison between the experimental tumor growth data and model predictions, and **Fig. 3B** shows the immune profile comparison. To validate the model, all parameters were kept the same between treatment groups except for the transmigration of immune cells in the tumor (for both tumor types), and the killing potential of T cells (for the GSC005 experiments); these parameters were adjusted in groups receiving anti-VEGF. These parameters were assumed to be affected by anti-VEGF treatment due to the improved vasculature and immune reprogramming of the TME caused by vascular normalization. The estimation of parameters is described in more detail in the Materials and Methods sub-section “Estimation of Model Parameters”, and the parameters are provided in Supplementary Table S2.

**Figure 3.**
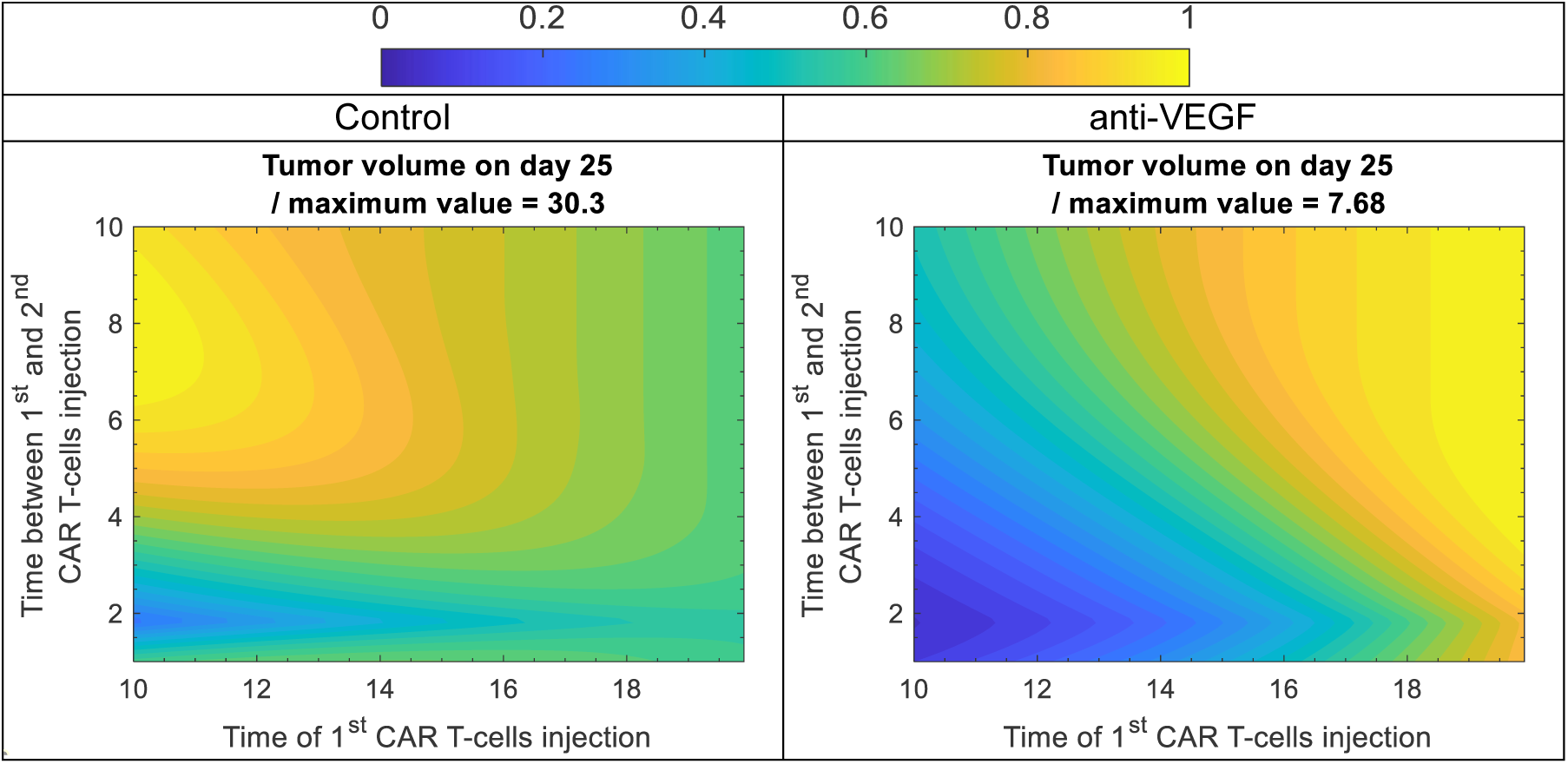
Tumor volume on day 25 for control and anti-VEGF with two systemic doses of CAR-T cells at different time of the 1st injection and different time between 1st and 2nd injection. The color scale from 0 to 1 corresponds to minimum and maximum value of tumor volume, i.e. tumor volume was divided with its maximum value to have a range between 0 to 1. The contour plots were generated by running simulations with two systemic doses of 106 CAR-T cells/cm3 of mouse vascular volume which is below the therapeutic threshold (see Fig. 4). The time of the 1st injection varies from day 10 to 20, and the time between 1st and 2nd injection varies from 1 to 10 days.

Anti-VEGF treatment of MGG8 tumors caused a reduction in tumor volume (**Fig. 3A**) combined with a reduction in the percentage of type 2 macrophages and an increase in the percentage of type 1 macrophages (**Fig. 3B** MGG8). The model accurately captures the changes in both tumor growth (*R*^2^ = 0.83) and macrophages (average relative difference 0.003). In the case of GSC005, the tumor growth curves of tumors that received CAR-T cell monotherapy are almost identical to control tumors, with no significant change in tumor volume. On the other hand, anti-VEGF treatment reduced the tumor volume, and the combination with CAR-T cell therapy (anti-VEGF + CAR-T) caused further reduction in tumor volume in both experimental and simulated data (**Fig. 3A** GSC005). The reduction of tumor growth with anti-VEGF was associated with increased infiltration of CD8^+^ effector T cells and CAR-T cells and reduced infiltration of immunosuppressive regulatory T cells (FOXP3^+^CD4^+^T cells, **Fig. 3B**). In this case, the model is also in good agreement with both tumor growth (*R*^2^ = 0.68) and immune profile data (average relative difference 0.17). In summary, the model was validated with the available in vivo data from two GBM cancer cell lines, and its results are in good agreement with the experimental data.

Next, we also validate our model with other data from the literature (24, 39) (see Supplementary Figure 1). Then we used our validated model to simulate various therapeutic scenarios and treatment protocols for combining CAR-T cell therapy and vascular normalization.

### Optimum dose and time of CAR-T cells injection

Then, we tested different systemic doses of CAR-T cells at different times of injection. Using the model, we performed analyses for a normalized vasculature (with anti-VEGF) and an abnormal vasculature (Control). Again, the only model parameters adjusted between the two cases were the transmigration of immune cells in the tumor and the killing potential of T cells, consistent with previous model calibration. On the day of CAR-T cells injection, the vasculature was assumed to be normalized, i.e., the time to reach vascular normalization was not considered. The dose of CAR-T cells varied from 10^3^ to 10^9^ cells/cm^3^ of mouse vascular volume (CAR-T cell concentration at the time of injection in mouse vascular volume, the volume of the circulating blood), and the time of injection varied from day 10 to day 20. Based on the published data, 2×10^6^ CAR-T cells were injected per mouse (25), giving a blood concentration of ∼1×10^6^ cells/cm^3^ after the division by the mouse vascular volume (29, 30). Here, the results are presented in terms of CAR-T cell concentration in the circulating blood at the time of injection, and the total number of CAR-T cells per mouse can be calculated by multiplying the concentration by the mouse vascular volume (38, 39). Fig. 4 shows the tumor volume and CD8^+^ T cells in the tumor compartment on day 25 for different combinations of doses and day of injection. Note that each quantity was divided by its maximum value to keep the range of each quantity from 0 to 1 and have the same colorbar. The maximum value of each quantity is provided in the title above each plot of Fig. 4.

**Figure 4.**
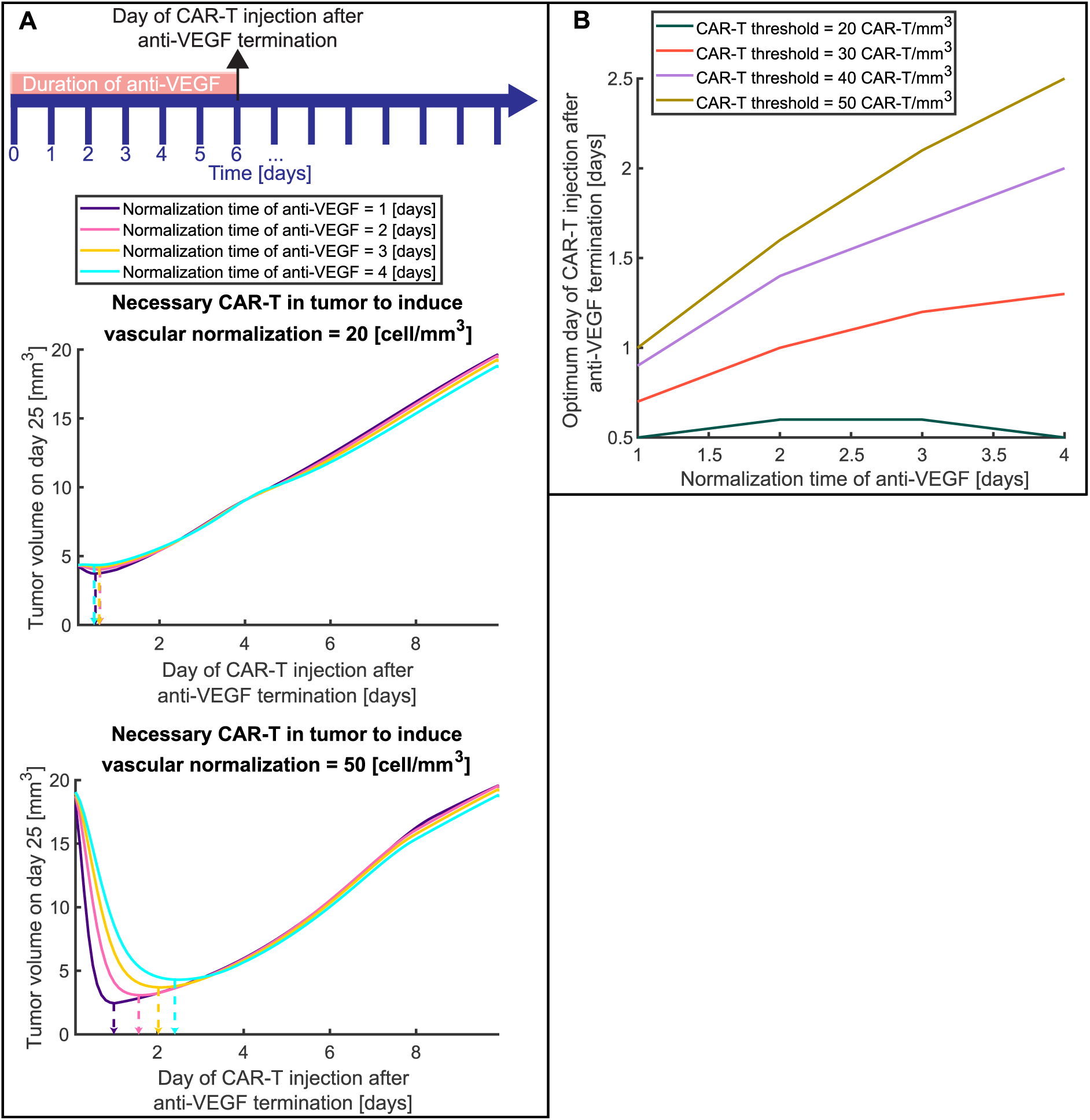
The optimal timing of systemic injection of CAR-T cells that produce anti-VEGF after anti-VEGF antibody normalization. A) Tumor volume on day 25 as a function of injection day after anti-VEGF termination for different normalization times of anti-VEGF. The timeline presents an indicative example of treatment protocol for the duration of anti-VEGF treatment and the day of CAR-T injection; the protocol varies, and the day of anti-VEGF termination and injection of CAR-T cells is the horizontal axis of Fig. 6A. The top plot presents such curves for the number of CAR-T cells that are needed to induce vasculature normalization in a tumor equal to 20 cells/mm3, and the bottom plot presents such curves for 50 cells/mm3. The vertical arrows with dashed lines indicate the optimum day of CAR-T cells injection where the tumor volume is minimum for each normalization time. B) The optimum day of CAR-T cell injection after anti-VEGF termination for different normalization times with anti-VEGF and different numbers of CAR-T cells per unit volume in the tumor that is needed to induce vasculature normalization (CAR-T threshold).

The results indicate that the effective dose to reduce tumor volume for the control groups is above 10^7^ whereas for the anti-VEGF groups is an order of magnitude lower, i.e, 2×10^6^ CAR-T cells (/cm^3^ of mouse vascular volume). Additionally, vessel normalization with anti-VEGF has a superior effect, with about half the maximum tumor volume on day 25: the maximum tumor volume for the Control is 28.6mm^3^ and for tumors treated with anti-VEGF is 13.15mm^3^. Furthermore, the normalized vasculature with anti-VEGF led to about a 5-fold higher maximum number of CD8^+^ T cells in the tumor: the number of CD8^+^ T cells/mm^3^ is 59 and 309 for Control and anti-VEGF treated tumors, respectively, indicating that vessel normalization also increases the accumulation of CD8+ T cells in the TME. The results for CD8^+^ T cells within the tumor (**Fig. 4**, left side) show that, above the therapeutic threshold of CAR-T cells dose, delaying the time of injection leads to a reduced number of CD8^+^ T cells in the tumor. This suggests that, above the therapeutic threshold of CAR-T cell dose, earlier CAR-T cell administration results in greater CD8^+^ T cell accumulation in the tumor and improved therapeutic outcomes.

There is also an optimum dose of CAR-T cells that simultaneously maximizes the number of CD8^+^T cells in the tumor (yellow color on the left plots of Fig. 4) and minimizes the tumor volume (blue color on the right plots of Fig. 4). This value is around 10^8^ and 10^7^ CAR-T cells (/cm^3^ of mouse vascular volume) for the control and anti-VEGF groups, respectively. When CAR-T doses fall below the therapeutic threshold, tumor response can become non-monotonic (producing the counter-intuitive dips) —for example, administering 10^5^–10^6^ cells (in the control case) early might yield weaker responses than slightly higher or lower doses or later injections. This arises from the interaction between innate and adaptive immunity and the available tumor antigen generated from the CAR-T cells. In these cases, the dose is insufficient, limiting antigen presentation and failing to sufficiently expand CD8⁺ T cells.

Overall, when the vasculature is normalized with anti-VEGF treatment, CAR-T cell therapy can be more effective with a 10-fold lower dose than the control. Also, normalization results in a better adaptive immune response with higher numbers of CD8^+^T cells, which in turn improves the therapeutic outcome and can potentially reduce side effects. Finally, when the CAR-T cell dose is above the therapeutic threshold, injecting CAR-T cells as soon as possible after anti-VEGF treatment leads to the best response.

### Optimum time interval between two systemic CAR-T cells injections

Following the investigation of the doses and timing of CAR-T cell therapy, we explored the administration of multiple doses of CAR-T cell therapy as another possible strategy to improve efficacy (40). Again, the analysis was performed for a normalized vasculature (with anti-VEGF) and an abnormal vasculature (control). The dose of CAR-T cells was set to 10^6^ CAR-T cells (/cm^3^ of mouse vascular volume), which is below the therapeutic threshold (see Fig. 4). This scenario tests whether an improved outcome can be achieved with lower, but multiple, doses, which can also reduce side effects. The time of the 1st injection varied from day 10 to 20, and the time between the 1st and 2nd injection varied from 1 to 10 days. Tumor volume on day 25 for control and anti-VEGF with two systemic doses of CAR-T cells is presented in **Fig. 5** for different times of the 1st injection and different time intervals between the 1st and 2nd injection. It must be noted that the second dose administration extends beyond day 25, so its full effect is not captured in **Figure 4**; however, these cases fall well above the optimal window, and thus, we use the same day for tumor volume representation as **Figure 3**.

**Figure 5.**
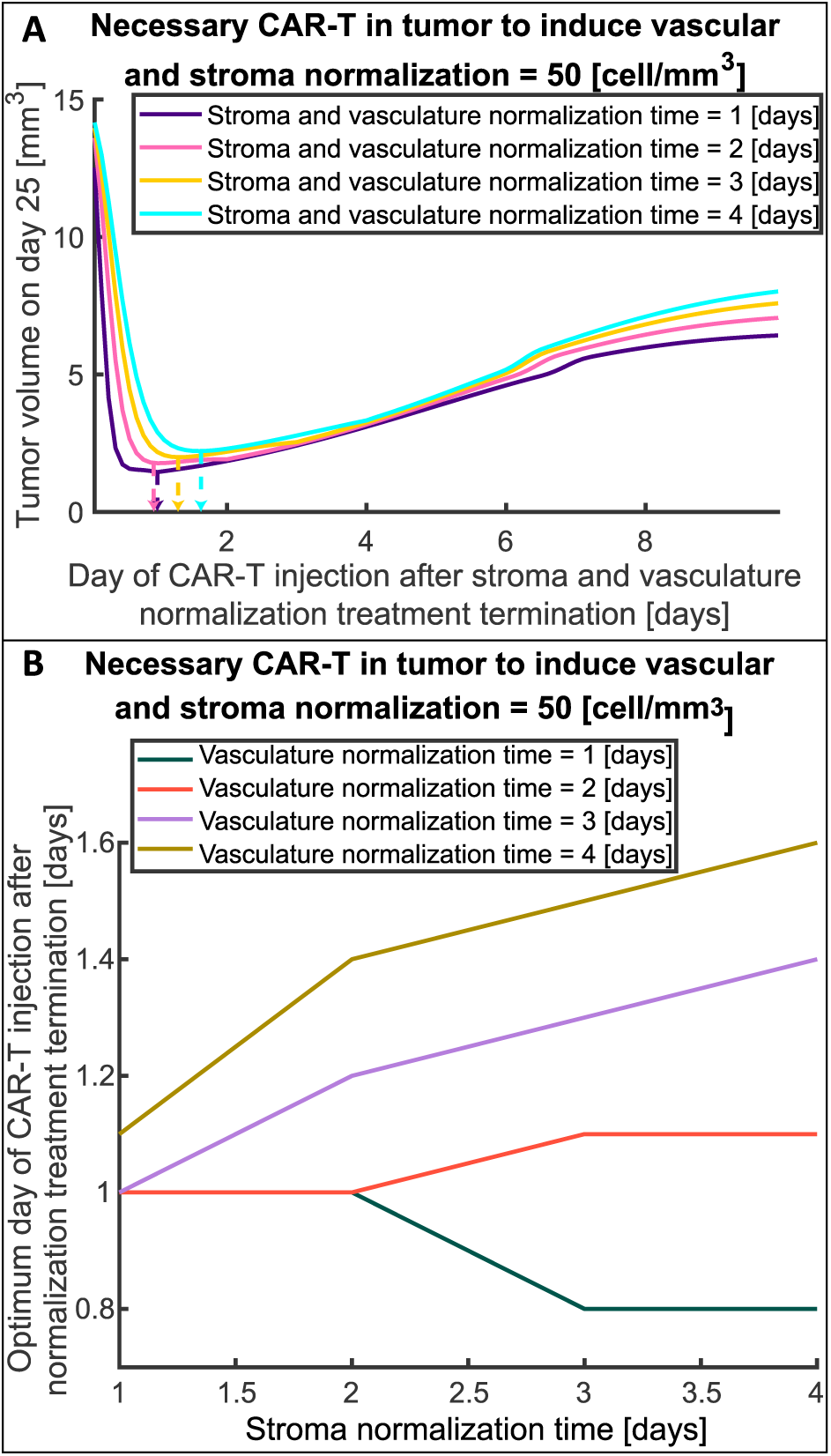
The optimal timing of systemic injection of FAP-CAR-T cells that produce anti-VEGF after treatment with stroma and vasculature normalizing agents. A) Tumor volume on day 25 as a function of injection day after stroma and vasculature normalization treatment termination for different stroma and vasculature normalization times (i.e., stroma and vascular normalization time equal to 1, 2, 3, or 4 days). The figure presents such curves for the number of CAR-T cells needed to induce vasculature and stroma normalization in the tumor, equal to 50 cells/mm3. The vertical arrows with dashed lines indicate the optimum day of CAR-T cells injection where the tumor volume is minimum for each normalization time. B) The optimum day of CAR-T cell injection after stroma and vasculature normalization treatment termination as a function of stroma normalization time for different vascular normalization times and number of CAR-T cells per unit volume in tumor needed to induce vasculature and stroma normalization equal to 50 cell/mm3 (CAR-T threshold).

In the case of the abnormal vasculature (control, left plot in **Fig. 5**), the distribution of the tumor volume is more uniform, with a very sharp and small optimal window indicated by the light blue color. Examining this in more detail, we see that injecting the first dose as soon as possible (day 10), requires the second dose to be injected between day 1.5 and 2 to achieve the lowest tumor volume. However, even with optimal time of injections, the tumor volume is quite large (about 10 mm^3^), and might need additional treatments. The dose of 10^6^ CAR-T cells (/cm^3^ of mouse vascular volume) is one order of magnitude below the therapeutic threshold for the control case, and even with two doses, the response is not sufficient.

In the case of a normalized vasculature (anti-VEGF, right plot in **Fig. 5**), the distribution of tumor volumes is more heterogeneous, with a larger optimal window indicated with the deep blue color, where the tumor volume is close to zero. In this case, if the first dose is injected as soon as possible (day 10), the second dose can be injected between day 1 and 2.75 without any significant change in the outcome. But if the 1^st^ dose is delayed for about 1 day (day 11), the window of the second injection becomes smaller, and the second dose needs to be injected between day 1.5 and 2.25 to achieve the same outcome. Furthermore, if the first dose is delayed more than a day, the window for the second dose becomes smaller and smaller until it closes, and when it closes, the tumor volume will be higher than the optimal case.

These results show that, when administered within the optimal window, two doses of CAR-T cells at a sub-therapeutic concentration threshold (see **Fig. 5**) can achieve similar results as a single dose with a concentration above the therapeutic threshold. An optimal window also exists for the non-normalized vasculature, but the treatment is not sufficiently effective, and more treatments are needed to further reduce the tumor volume. Anti-VEGF prolongs the optimal window, and, in combination with CAR-T cell injections, reduces tumor volume. In both cases, when the first dose is injected as early as possible, the margin of improvement with a second dose is larger due to the larger optimal window of the second injection. Additionally, the best time for the second dose is about 1.5 days after the first dose, which might be related to the maximum time of CAR-T cell proliferation under antigen stimulation, which was estimated around this value during the calibration of the model. Due to the continuous antigen stimulation, the CAR-T cells from the first dose become exhausted and can no longer proliferate. As a result, the second dose introduces fresh CAR-T cells capable of proliferating in response to ongoing antigen stimulation, thereby expanding the CAR-T cell population within the tumor and ultimately enhancing treatment efficacy.

### Optimum time of injection of anti-VEGF–producing CAR-T cells after vascular normalization

We next investigated whether pre-induction of vascular normalization using anti-VEGF, followed by administration of anti-VEGF-producing CAR-T cells, could improve outcome. Such CAR-T cells can simultaneously induce cancer cell cytolysis and maintain normalization of tumor vessels (41). We hypothesize, however, that even though these engineered T cells secrete anti-VEGF themselves, pre-conditioning the tumor vasculature with anti-VEGF may still be required to ensure efficient homing and infiltration of the CAR-T cells into the TME. CAR-T cells will be able to induce vascular normalization by anti-VEGF production only when a critical number of CAR-T cells have entered the TME.

To find the optimum treatment schedule with anti-VEGF antibodies before the systemic injection of anti-VEGF-secreting CAR-T cells, we explored two crucial factors. The first was the normalization time, defined as the duration of antibody treatment needed to restore adequate vessel perfusion, prune abnormal vessels, and increase immune cell adhesion. The second was the threshold of CAR-T cells density within the tumor—i.e., the minimum number of CAR-T cells per unit volume required to secrete sufficient anti-VEGF locally to maintain or further the normalization state originally induced by the monoclonal antibody. This second parameter dictates how many CAR-T cells must infiltrate the tumor and depends on the ability of CAR-T cells to produce anti-VEGF. To test these factors, we varied the pre-conditioning normalization time of anti-VEGF from 1 to 4 days and the necessary CAR-T cells density from 20 to 50 [cells/mm^3^]. In the model, the transmigration of immune cells into the tumor and the killing potential of T cells, which were assumed to be affected by vascular normalization, were transiently varied from the control to the normalized state, either following anti-VEGF antibody treatment or when CAR-T cell density exceeded the normalization threshold. The administration of anti-VEGF always starts at the same time (day 0 of CAR-T cells injection after anti-VEGF termination in **Fig. 6A**) and stops on the day of CAR-T cell injection. The normalization time is not the duration of anti-VEGF administration, but rather the duration of antibody treatment needed to fully restore the vessels. That is, for a specific normalization time, the duration of anti-VEGF treatment might be shorter or longer than the normalization time; when the anti-VEGF treatment is terminated, CAR-T cells are injected. The timeline in **Fig. 6A** is an example that represents exactly these two time points: the duration of anti-VEGF treatment (6 days) and the time of CAR-T injection (at day 6), which varies and is indicated by the horizontal axis of **Fig. 6A**. The goal is to identify the optimal scheduling for combining anti-VEGF antibody and anti-VEGF-producing CAR-T cell therapy by determining how the timing of these two factors induces CAR-T cell infiltration and sustained vascular normalization.

**Figure 6:**
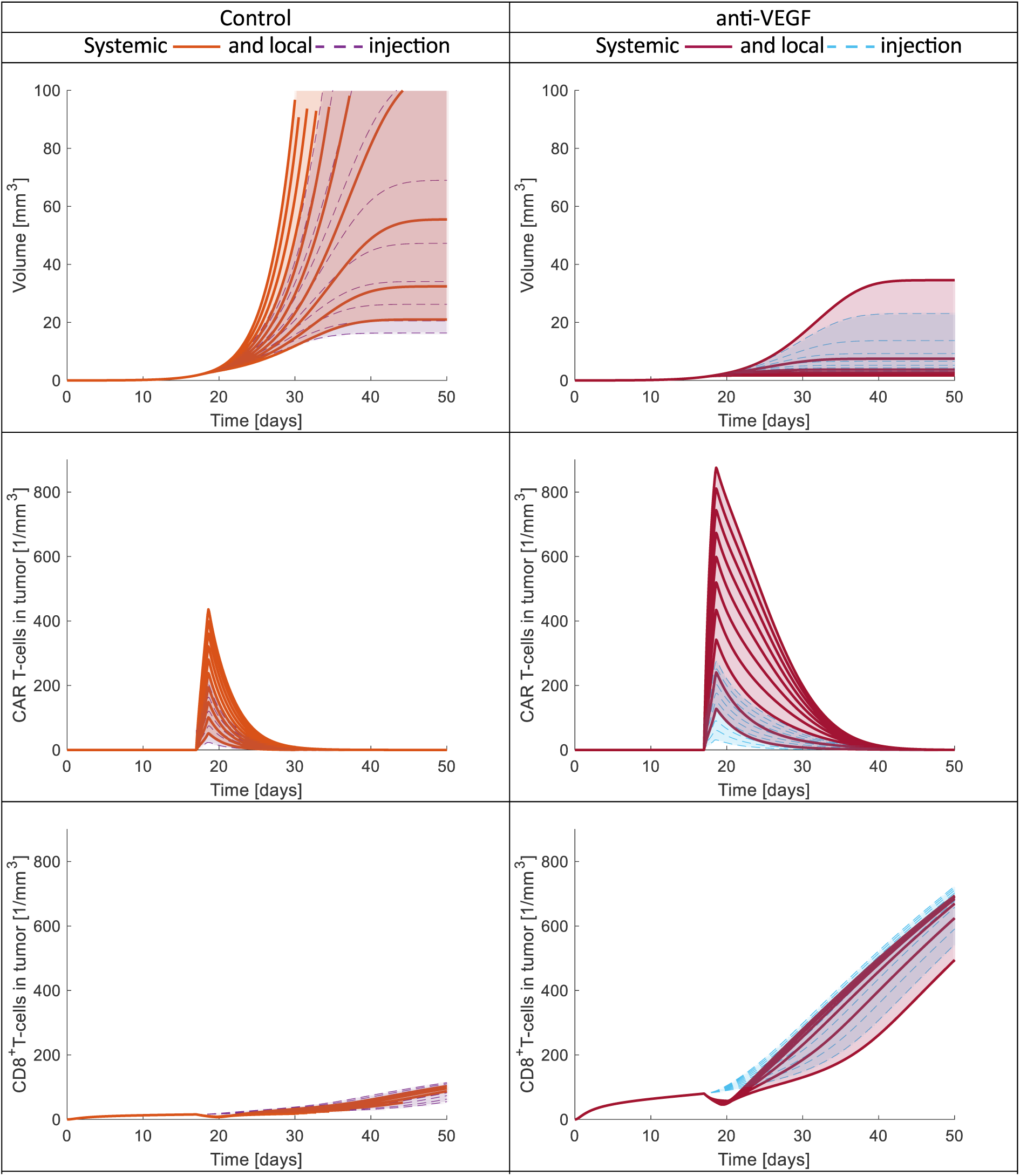
Time evolution of tumor volume, and CAR-T cells and CD8+ T cells in the tumor for different CAR-T cell doses administrated systemically or locally. On the left are the results for the control (abnormal vasculature) and on the right for anti-VEGF (normalized vasculature). The first, second and third row presents the tumor volume, CAR-T cells in tumor and CD8+T cells in tumor respectively. The different curves are from different doses. For the systemic injection the dose ranges from 106 to 107 cells/cm3 of mouse vascular volume and for the local injection from 10 to 100 cells injected into the tumor (independent of tumor volume).

**Fig. 6A** presents the tumor volume on day 25 as a function of the day of CAR-T cell injection after termination of anti-VEGF treatment for different normalization times of anti-VEGF antibodies and two values of the necessary number of CAR-T cells needed to induce vascular normalization (**Fig. 6A** top plot 20 cells/mm^3^ and bottom plot 50 cells/mm^3^). In cases where few CAR-T cells are needed to infiltrate the tumor in order to induce vascular normalization (**Fig. 6A**, value of 20 cells/mm^3^), no significant improvement was observed by introducing anti-VEGF pre-treatment. The dashed arrows (**Fig. 6A**) indicate the possible minimum volume that can be achieved by injecting the CAR-T cells at the optimum time (about 0.5 day after anti-VEGF). For this case, the minimum tumor volume is approximately the same whether CAR-T cells are injected at about the same time with anti-VEGF or at the optimum time. On the other hand, when higher values of CAR-T cells are needed to induce vascular normalization (**Fig. 6A**, value of 50 cells/mm^3^), a significant reduction of tumor volume can be achieved by using anti-VEGF treatment first and then CAR-T cell therapy. In this case, the possible minimum volume [dashed arrows (**Fig. 6A**)] is 10 to 4 times lower than the maximum tumor volume (worst case), which is achieved when CAR-T cells are injected at about the same time with anti-VEGF (first point of **Fig. 6A** with value of 50 cells/mm^3^) or after 10 days (last point of **Fig. 6A** with value of 50 cells/mm^3^). The reduction of tumor volume depends on the normalization time with anti-VEGF. Specifically, when more time is needed to induce vascular normalization with anti-VEGF, more anti-VEGF pre-conditioning treatment is needed, which delays CAR-T cell therapy and thus leads to larger tumor volumes. Overall, for all cases, there is an optimum time for anti-VEGF termination and injection of CAR-T cells.

Titration of the timing of CAR-T cell injection after anti-VEGF termination is presented in **Fig. 6B**. When low values of CAR-T cells are needed to induce vascular normalization (CAR-T threshold 20 cells/mm^3^) the optimum timing of CAR-T cell injection is nearly independent of normalization time and equal to 0.5 [day] (about half day after anti-VEGF). But when the CAR-T cell threshold increases, the dependence on normalization time also increases (see Fig. 6B, and curves with CAR-T threshold 30, 40 and 50 cells/mm^3^). Interestingly, the optimum day of CAR-T cell injection after anti-VEGF termination differs from the normalization time, and there is an intermediate point between anti-VEGF initiation and normalization time where the benefits of vascular normalization balance the negative effect of tumor progression. For example, when the CAR-T threshold is 50 cells/mm^3^ and the normalization time is 4 days, the optimum time of CAR-T cell injection after anti-VEGF termination is about 2.5 days. At the optimum time, anti-VEGF treatment induces sufficient vascular normalization to ensure that enough CAR-T cells can infiltrate the tumor to maintain the normalization and simultaneously exert cytotoxic effects.

### Optimum time of injection of anti-VEGF–producing FAP-CAR-T cells after vascular and stroma normalization

In desmoplastic tumors, such as subtypes of pancreatic and breast tumors and sarcomas, blood vessels are compressed and perfusion is compromised. Thus, treatments that only affect vessel permeability will not improve perfusion, as the vessels remain compressed (27). In these cases, vascular normalization can be synergistically combined with stroma normalization, as previous studies suggest (27, 28) to optimize therapeutic outcomes in specific types of solid tumors (**Fig. 1**). Vascular and stroma normalization can be achieved by combining anti-VEGF treatment with stroma normalizing agents (e.g. Losartan (42) and Ketotifen(43)) to pre-condition the tumor microenvironment prior to therapy. There are also CAR-T cells that can be engineered to simultaneously target not only cancer cells but also cancer associated fibroblast, known as FAP (Fibroblast Activated Protein)-CAR-T cells (44, 45), and FAP-CAR-T cells that also produce anti-VEGF.

For this scenario, we investigated whether the combination of vascular and stroma normalization can improve the outcome of cancers treated with anti-VEGF-producing FAP-CAR-T cells. Such CAR-T cells can simultaneously target cancer cells and cancer-associated fibroblasts (CAFs) that express Fibroblast Activated Protein (44, 45). Using our model, we tested whether pre-conditioning the tumor with vascular and stroma normalizing agents before systemic injection of anti-VEGF producing FAP-CAR-T cells can further improve the treatment outcome. It was assumed that stroma normalization induced either by stroma normalizing agents or FAP-CAR-T cells causes a 50% increase in the vascular volume of the tumor. Vascular normalization with anti-VEGF and anti-VEGF producing CAR-T cells was considered as described in the previous section.

**Fig. 7A** presents the tumor volume on day 25 as a function of the day of CAR-T cell injection after termination of stroma and vasculature normalization treatment for different normalization times of these agents. The horizontal axis of **Fig. 7A** indicates the duration of normalizing agents treatment and the time of CAR-T injection similar to **Fig. 6A** and the indicative timeline. Although the time required for vascular and stroma normalization could be different, here we assumed that they are the same (only for representation in **Fig. 7A**). The necessary number of CAR-T cells that are needed to induce vascular and stroma normalization (i.e., CAR-T threshold in **Fig. 7A**) was set to 50 cells/mm^3^ which was found to have the highest potential for improved therapy in previous results (**Fig. 6**). The combination of stroma and vasculature normalizing agents and anti-VEGF producing FAP-CAR-T cells results in lower tumor volume on day 25 (**Fig. 7A**) in comparison to anti-VEGF and anti-VEGF producing CAR-T cells (**Fig. 6A**, 50 cells/mm^3^), and results in a more stable response after the optimum time of injection (after dashed arrows in **Fig. 7A**). The combination of stroma normalization + vascular normalization + anti-VEGF producing FAP-CAR-T cells can potentially improve survival, even if the CAR-T cells are injected after the optimum time point (**Fig. 7A**) because the tumor volume does not increase as rapidly as in the case of anti-VEGF + anti-VEGF producing CAR-T cells (**Fig. 6A**). At the optimum time of injection (indicated by dashed arrows), the tumor volume is reduced 8.5 to 6.5 times in comparison to the maximum tumor volume (worst case), which is achieved when CAR-T cells are injected at about the same time as stroma and vasculature normalizing agents (first point of **Fig. 7A**). The current treatment with combined stroma and vasculature normalizing agents with anti-VEGF producing FAP-CAR-T cells has a shorter optimum time of CAR-T cell injection than the previous case of combined anti-VEGF + anti-VEGF producing CAR-T cells. This optimum time ranges from 0.8 to 1.6 days (**Fig. 7B**) for the different normalization times of stroma and vasculature normalizing agents; the lower values compared to **Fig. 6B** can be explained by the synergistic effects of vascular and stroma normalization, which increase the infiltration of CAR T cells, thus achieving the CAR-T threshold sooner than if only vascular normalization is considered.

Optimization of the timing of CAR-T cell injection after termination of stroma and vasculature normalization treatment is presented in **Fig. 7B** for different stroma and vasculature normalization times. For shorter vascular normalization times (1 and 2 days), the optimum day of CAR-T cell injection is around day 1 and ranges from 0.8 day (about 17 hours) to 1.1 day (about 26 hours), and does not show a dependence on stroma normalization time. On the other hand, for longer vascular normalization times (3 and 4 days) the optimum day of injection depends on stroma normalization time and ranges from day 1 to day 1.5. Again, the optimum day of CAR-T cell injection after stromal and vascular normalization treatment termination in most cases is not the same as normalization times, and there is an intermediate point between the initiation of stroma and vasculature normalizing agents and normalization times where the benefits of vascular and stroma normalization balance the negative effect of tumor progression. Overall, when both vascular and stroma normalization are included, outcomes are better and responses are more stable, even when the CAR-T cells are injected after the optimum time.

### Local CAR-T cell injection outperforms systemic delivery

Given that in brain tumors such as GBM, the CAR-T cells can be injected locally, intracranially or intraventricularly, (46, 47), we also simulated the local injection of CAR-T cells. **Fig. 8** presents the time evolution of tumor volume as well as the concentrations of CAR-T cells and CD8^+^ T cells in the tumor. We compare local to systemic injections for different doses of CAR-T cells. For the analysis, we performed simulations for both control (abnormal vasculature) and anti-VEGF cases (normalized vasculature). It is important to note that in **Fig. 8**, for the local injection, the number of CAR-T cells (absolute value) was set 5 orders lower than the number of CAR-T cells /cm^3^ of mouse vascular volume injected systemically. The range of the number of CAR-T cells for local injections was chosen to have similar effects on tumor volume as the systemic injections.

The tumor volume for both local and systemic injections has a similar range in control or anti-VEGF but the maximum number of CAR-T cells in the tumor is lower in the local injection compared to systemic one [see tumor volume and number of CAR-T cells in **Fig. 8**]. These findings come from the fact that the local injection is more targeted because CAR-T cells are immediately injected into the TME and proliferate at the same time as they cytolyze cancer cells; thus, lower dose and lower amount of CAR-T cells in tumor can control the tumor volume. For both systemic and local injections, the normalized vasculature has better response than the non-normalized one due to the immunostimulatory environment which allows the activation and infiltration of more CD8^+^ T cells (see number of CD8^+^ T cells in **Fig. 8**). The local injection of CAR-T cells also increases the CD8^+^ T cells in the tumor because the systemic injection restricts the transport of CD8^+^ T cells in the blood circulation as it indicated by the slight decrease of CD8^+^ T cells at the time of CAR-T cells injection in both Control and anti-VEGF (see number of CD8^+^ T cells in **Fig. 8**).

In general, the local injections have many advantages in comparison with systemic injections. Firstly, assuming optimal doses, a local injection can have the same effect as a systemic injection, but with fewer injected cells, thus minimizing side effects. Secondly, the CAR-T cells are immediately inside or in close contact with the TME, avoiding barriers to transport through the blood circulation and into the tumor tissue. Thirdly, local injections do not hinder the circulation of other immune cells such as CD8^+^ T cells, and thus, more CD8^+^ T cells can infiltrate the tumor and improve treatment response. Finally, we found that the normalized vasculature results in the best outcome for both local and systemic injections.

## Discussion

In this study, we developed a PBPK model that simulates the behavior of endogenous immune cells and CAR-T cells in the treatment of GBM using a combination of anti-VEGF and CAR-T cells therapies. The model was validated with both tumor growth data and immune profile data from two different cancer cell lines (25, 38). A generic optimization algorithm was used to estimate the unknown parameters governing the model from which only key parameters such as the transmigration of immune cells and killing potential of T cells were assumed to be affected by vascular normalization and anti-VEGF due to improved normalized vasculature and the reprogramming of the TME from immunosuppressive to immunosupportive. Anti-VEGF normalization increases the infiltration of CD8^+^ effector T cells (25), M1 (38), CAR-T cells (24, 25) and reduces the infiltration of M2 (38) and Tregs (25). Vessel normalization induces immune reprogramming within the TME, which favors the treatment of solid tumors and is evidenced by both simulated and experimental results. Subsequently, we used the model to test different treatment scenarios and identify optimum treatment strategies for the combination of vasculature and stroma normalizing agents and CAR-T cells or anti-VEGF producing CAR-T cells, or anti-VEGF producing FAP-CAR-T cells therapies for GBM and other solid tumors. Developing such a mathematical model offers substantial advantages, particularly in the context of complex treatment strategies where experimental testing of every possible variable combination is impractical. For example, exploring the effects of variations in CAR-T cells dose, timing of administration, injection route, or coordination with vasculature and stroma normalizing agents would require extensive in vivo experimentation that is often limited by cost, time, and ethical considerations. Thus our PBPK model provides a powerful in silico platform to systematically explore these scenarios, prioritize promising strategies, and refine experimental designs before moving to preclinical or clinical stages.

The present study underscores the critical interplay between TME normalization achieved through vascular normalization (anti-VEGF) therapy, stroma normalization and the efficacy of CAR T cell treatment. A primary observation is that the normalization of tumor vasculature can dramatically enhance CAR-T cell infiltration when they injected systemically, thereby allowing effective therapeutic outcomes with a CAR-T cell dose that is an order of magnitude lower than that required in the absence of normalization. This finding is significant because it not only suggests a method to boost the therapeutic efficacy of CAR-T cell therapy but also offers the possibility to reduce side effects associated with higher cell doses. Furthermore, the normalized vasculature facilitates a robust adaptive immune response, as evidenced by the observed increase in CD8⁺ T cell infiltration. This infiltration is crucial because CD8⁺ cytotoxic T lymphocytes contribute to sustained antitumor activity. Another key insight from the results is the importance of timing in the systemic injection of CAR-T cells. When CAR-T cells dose exceeds a certain therapeutic threshold, early injection of cells leads to superior outcomes with lower tumor volume and higher levels of CD8⁺ effector T cells.

The analysis also reveals that there is an optimal window for the timing of two sequential systemic injections. Administering two systemic doses of CAR-T cells at subtherapeutic threshold levels— provided they are timed within the optimal window—yielded outcomes comparable to a single high dose when the vasculature is normalized with anti-VEGF. More specifically, the optimal time window of the 2^nd^ dose closes and becomes smaller if the first dose is delayed; thus, treatment is better when started as early as possible. The optimum time interval between doses is approximately the maximum period of CAR-T cells proliferation under antigen stimulation. By the time of the second dose, the initial CAR-T cells population begins to exhibit signs of exhaustion, and the introduction of fresh CAR-T cells helps sustain an expanding antitumor response. The synchronization of dosing with this proliferation dynamic is a critical factor that appears to mitigate T cell exhaustion and lead to optimum treatment.

In the case of CAR-T cells that produce anti-VEGF, normalization with anti-VEGF before CAR-T cells systemic injection might improve the outcome under certain conditions. The conditions depend on the normalization time of the vasculature with anti-VEGF and the number of CAR-T cells in the tumor required for vascular normalization. The normalization time depends on the tumor type, the abnormality level, and the treatment dose, while the number of CAR-T cells in the tumor needed to induce vascular normalization also depends on their ability to produce anti-VEGF. The optimum time of CAR-T cell administration after anti-VEGF termination is an intermediate point between anti-VEGF initiation and the normalization time, and increases as more CAR-T cells are needed to induce vascular normalization. This intermediate timing likely reflects a balance where the normalized vasculature sufficiently supports CAR-T cell infiltration, and tumor progression has not advanced to a point where it significantly compromises treatment efficacy.

The results demonstrate that the combined approach of vascular normalization and stroma normalization, followed by the administration of anti-VEGF producing FAP-CAR-T cells, significantly enhances therapeutic outcomes compared to combination of vascular normalization and anti-VEGF producing CAR-T cells. Specifically, the integration of both normalization strategies prior to CAR-T cell injection led to improved vasculature within the tumor and facilitated more effective immune cell infiltration. This synergistic normalization effect allowed the CAR-T cells to reach the normalization threshold more rapidly, thus achieving tumor volume reductions. Importantly, the combined treatment exhibited a more robust and stable tumor response even when anti-VEGF producing FAP-CAR-T cells were injected after the optimal time point, suggesting greater flexibility in clinical application. Furthermore, the analysis of optimum injection timing revealed that, for tumors with shorter vasculature normalization time, the optimal time of CAR-T cell administration remained relatively constant, whereas tumors with longer vasculature normalization time had more variability, and out efficacy was dependent on stroma normalization time. These findings underscore the importance of synchronized vascular and stroma normalization to enhance anti-VEGF producing FAP-CAR-T cells efficacy even though these CAR-T cells can induce stroma and vascular normalization by themselves.

Finally, local injections were found to have several benefits over systemic administration. Because local administration directly introduces CAR-T cells into or near the TME, it bypasses the common issue of cell loss due to circulatory dispersion, ensuring a higher local concentration of effector cells. This direct approach minimizes potential systemic side effects and may facilitate a more rapid and robust immune response. Moreover, by preserving the normal circulation of other immune cells, particularly CD8⁺ T cells, local injections might further fortify the antitumor immune milieu. Importantly, both administration routes achieve the best outcomes when supported by a normalized vascular environment, emphasizing the fundamental role of anti-VEGF treatment in this multi-modal therapeutic strategy. The current modeling strategy allows simulations of several treatment scenarios, and our analysis leads to optimum timing and doses that can boost future in vivo experiments.

## Materials and Methods

### Modeling strategy and testing scenarios

In this study, we developed and validated a PBPK model to simulate and predict the behavior of immune cells, including CAR-T cells, in the treatment of glioblastoma (GBM) using both anti-VEGF and CAR-T cells therapies. The modeling strategy integrated published in vivo experimental data (24, 25, 38, 39) to capture the dynamic behavior, and interactions of various immune cell populations with cancer cells. A generic optimization algorithm was used to estimate the unknown parameters governing the model. Then we used the model to test different doses of CAR-T cells injected systemically at different times of injection for cases that are normalized or not with anti-VEGF. Additionally, two systemic injections of CAR-T cells with different timing between doses and different time of initial injection were simulated for cases where vessels are normalized or not with anti-VEGF. Another testing scenario was to first induce vascular normalization using anti-VEGF and then inject systemically CAR-T cells that also produce anti-VEGF and can simultaneously induce cancer cell cytolysis and vascular normalization. The crucial parameters for this scenario were the normalization time, namely the time needed to normalize the vasculature with anti-VEGF and the number of CAR-T cells per unit volume needed to keep the vasculature normalized – specifically how many CAR-T cells are needed to produce enough anti-VEGF (this parameter also depends on the ability of CAR-T cells to produce anti-VEGF). These two parameters were varied along with the time of systemic CAR-T cells injection after anti-VEGF to find the best protocol and determine when pre-normalization with anti-VEGF is necessary before initiation of CAR-T cells therapy. We further extended our analysis be combining vascular and stroma normalizing agents and Fibroblast Activated Protein (FAP)-CAR-T cells (44, 45) that also produce anti-VEGF. Finally, local and systemic injections of CAR-T cells (48, 49) with different doses were tested.

### PBPK model of immune response and CAR-T cells therapy for solid tumors

The developed PBPK model is a comprehensive computational framework that simulates the biodistribution and interactions of immune cells across multiple physiological compartments. It incorporates 10 cell types, including Dendritic Cells (DCs), Macrophages type 1 and 2 (M1 and M2 respectively), regulatory T cells (Tregs), naïve and effector CD8^+^ T cells (Tn and TE1 respectively), Antigen-Presenting Cells (APCs), CAR-T cells (TE2), viable and dead tumor cells (Tv and Td respectively) from which 7 cell types (all excluding Tn, Tv and Td) are circulating throughout the compartments. The model spans 11 organ compartments (such as the Lungs, Liver, Gastrointestinal Tract, Spleen, Heart, Kidneys, Skin, Muscle, Bone, Lymph Nodes, and Tumor) along with blood and lymph circulatory systems, see **Fig. 2A**. Each organ is divided into vascular and extravascular sub-compartments that are interconnected through blood and lymphatic vessels, facilitating a realistic simulation of cell transport. The model employs a set of ordinary differential equations (ODEs) to ensure population conservation for each cell type. The population conservation equations track the accumulation of cells by incorporating cellular incoming and leaving fluxes, cellular proliferation/death, cytolysis, phagocytosis, activation against tumor antigen, deactivation induced by regulatory cells (Tregs and M2), and transport across sub-compartments. Additionally, in the vascular space, the freely circulating cells can transmigrate from vascular to extravascular space and from extravascular space cells can recirculate through lymphatic vessels. The population conservation equations for each cell type and each compartment consist of one equation in the vascular space and one in extravascular space. Special attention is given to the tumor compartment, where in the extravascular space, tumor cell proliferation and immune cell interactions, including cytolysis and phagocytosis, are modeled with the tumor volume being varied simultaneously based on the net change in cell numbers. Overall, the PBPK model consists of 164 ODEs and 1 algebraic equation (2 sub-compartments times 7 immune cell types times 11 compartments plus seven equations for the blood circulation of the 7 immune cells, two equations for the viable and dead tumor cells in tumor extravascular space, one extra equation for tumor growth and one algebraic equation for the Naïve CD8^+^ T cells in lymph node extravascular space) which are solved numerically using an implicit Euler method via COMSOL Multiphysics 5.6.

### Estimation of Model Parameters

To accurately capture the dynamics of immune-tumor interactions, a generic optimization algorithm was used to estimate the parameters governing the model. Many parameters were derived from a previous PBPK modeling publication (29), while the new parameters were estimated by fitting the model to experimental data, including tumor growth measurements, multiphoton microscopy and immunofluorescence data for CAR-T cells quantification, and flow cytometry data for immune cell profiling [effector CD8^+^ T cells, Tregs (FOXP3^+^CD4^+^T cells), CAR-T cells, M1 and M2] (25, 38). Parameter estimation was conducted for two distinct GBM cancer cell lines - GSC005 (murine) and MGG8 (human), see Supplementary Table 2 and 3.

We used an optimization algorithm to estimate tumor growth characteristics, immune response parameters, and immune cell trafficking rates. Anti-VEGF was assumed to affect two main processes: (i) immune cell infiltration into the tumor via improved, normalized vasculature, and (ii) T cell cytotoxic potential through reprogramming of the tumor microenvironment from immunosuppressive to immunosupportive. For the murine GBM model, 12 parameters, such as tumor growth rate, initial tumor volume, CAR-T proliferation dynamics, and immune suppression rates, remained constant across treatment groups, while 8 parameters related to immune cell transmigration and T cell killing capacity differed between anti-VEGF and control groups. In the human GBM model, conducted in immunodeficient mice, only parameters unrelated to adaptive immunity were retained. Seven parameters remained constant, and four transmigration-related parameters varied with anti-VEGF treatment. These calibrated parameters allowed us to simulate a range of therapeutic scenarios and identify optimal strategies for combining CAR-T therapy with anti-VEGF in GBM.

## Acknowledgments

We would like to thank Drs. Somin Lee, Vasiliki Slamet, Taylor Uccello and Xingjian Zhang for their helpful comments on this manuscript. This project was supported in part by funding from the European Research Council (ERC) under the European Union’s Horizon 2020 research and innovation program (grant agreement nos 863955, 101141357) to **T. Stylianopoulos**, grants R01CA247441, R21EB031982 and 5U01CA2618425 to **L.L. Munn**, grants R01-CA259253, R01-CA208205, R01-NS118929, U01-CA261842, and U01-CA 224348, Outstanding Investigator Award R35-CA197743 and grants from the National Foundation for Cancer Research, Jane’s Trust Foundation, Niles Albright Research Foundation and Harvard Ludwig Cancer Center to **R.K. Jain**.

---

**Figure 1.**
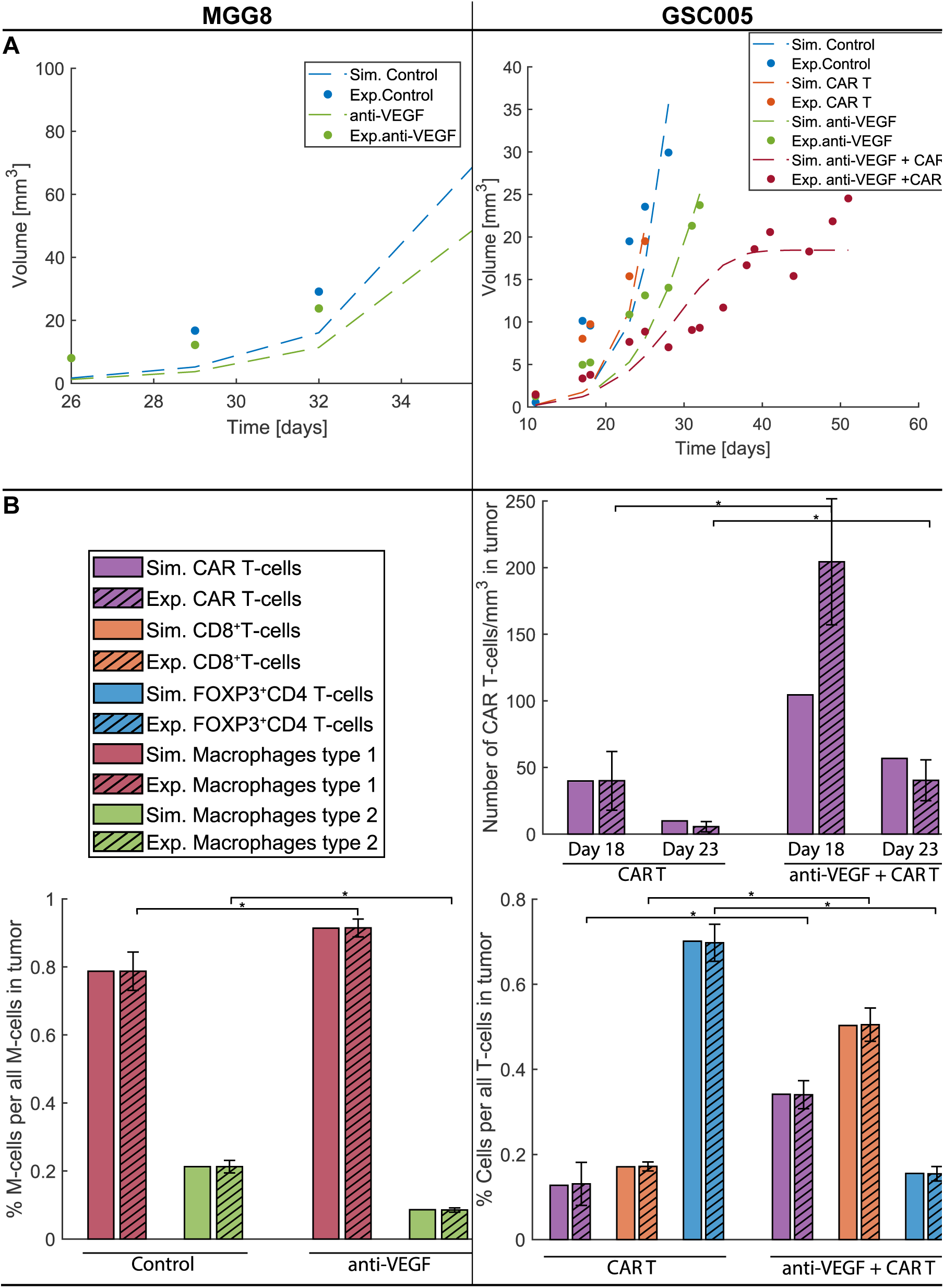
Comparison of model predictions with tumor size and immune profile data for two different GBM cell lines (MGG8 on the left and GSC005 on the right). A) Experimental tumor growth data (bullet points) and model predictions (continues dashed lines). B) Immune profile data for CAR-T cells, effector CD8+ T cells, FOXP3+CD4+ T cells (Tregs), macrophages type 1 (M1) and type 2 (M2), where the experimental data are presented with striped bars ±SEM, and the simulated data are presented with plain bars. * Experimental treatment groups with statistically significant different average values (25, 38).

**Figure 2.**
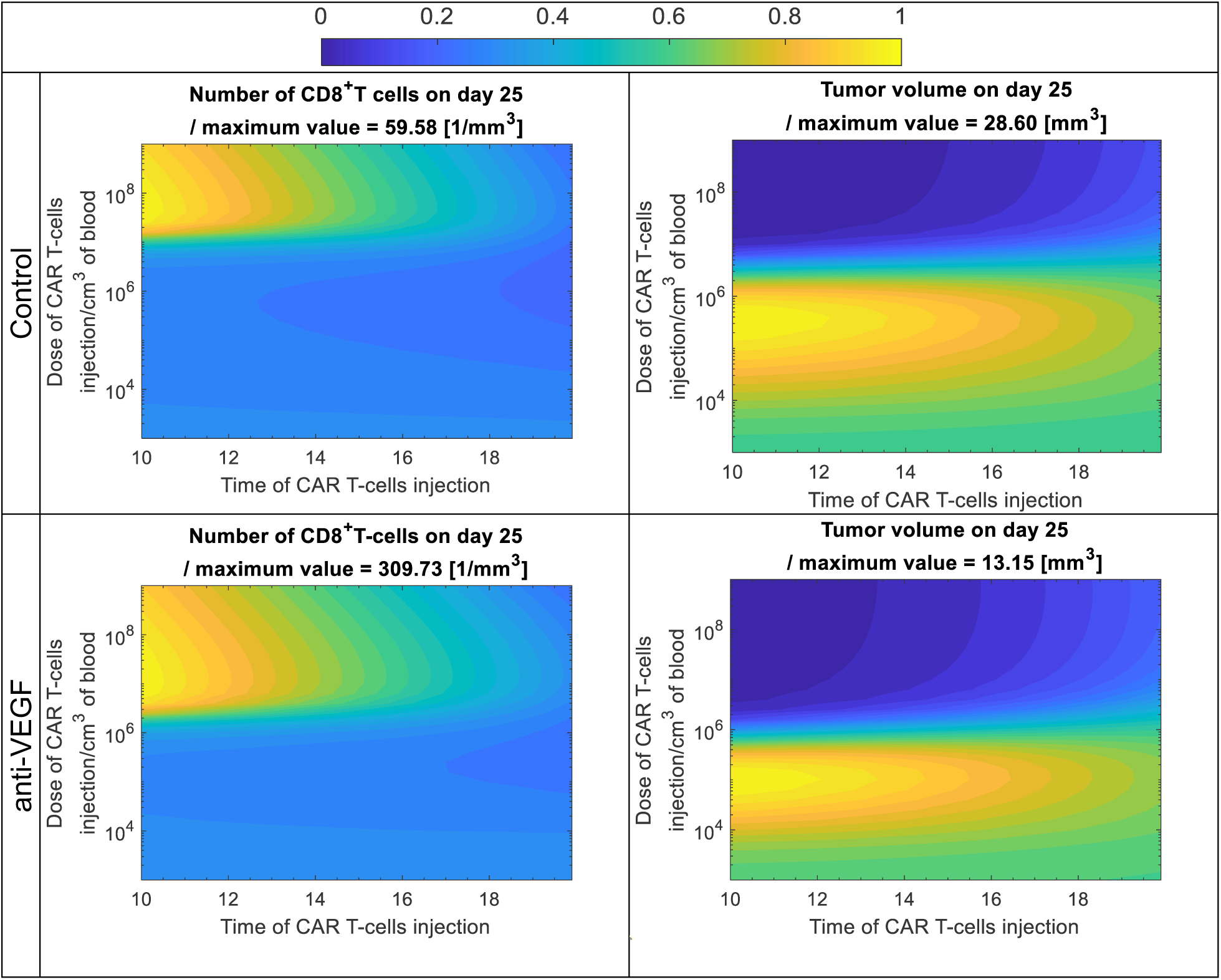
Tumor volume and CD8+ T cells in the tumor compartment on day 25 for control and anti-VEGF with different doses of CAR-T cells and different times of systemic injections. The color scale from 0 to 1 corresponds to minimum and maximum value of tumor volume (contour plots on the right) and CD8+ T cells in the tumor compartment (contour plots on the left), i.e. each quantity was divided with its maximum value to have a range between 0 to 1. The maximum value of each quantity is provided in the title above each contour plot. The contour plots were generated by running simulations with different systemic doses of CAR-T cells, varying from 103 to 109 / cm3 of mouse vascular volume (CAR-T cells concentration in mouse vascular volume at the time of injection), and different injection times, varying from day 10 to 20 after normalization of the vessels in anti-VEGF group.

## Notes

### Competing Interest Statement

R.K. Jain received consultant fees from SynDevRx; owns equity in Enlight and SynDevRx, and received grants from Sanofi. L.L. Munn receives equity from Bayer and is a consultant for SimBiosys. Neither any reagent nor any funding from any of the above organizations was used in this study. Other co-authors have no conflict of interest to declare.

