## Supplementary Information for "Physiologically-based pharmacokinetic model for CAR-T cells delivery and efficacy in solid tumors"

### EQUATIONS:

The physiologically based pharmacokinetics (PBPK) model was utilized to simulate the dynamic behavior and distribution of various immune cell populations across multiple organs (compartments). The model includes 10 different cell types: dendritic cells (DCs), macrophages type 1 (M1) and type 2 (M2), regulatory T cells (Tregs), naïve CD8+ T cells, viable and dead tumor cells (Tv and Td respectively) and naïve (Tn) and tumor specific CD8+ effector T cells (TE1), CAR-T cells (TE2) and antigen-presenting cells (APCs). The model tracks seven out of ten cell types (all excluding Tn, Tv and Td) as they are circulating throughout 11 key organs/compartments, including: lungs, liver, gastrointestinal tract, spleen, heart, kidneys, skin, muscle, bone, lymph nodes, and "tumor". Tn cells are calculated only in the lymph nodes where they exist in large quantities, and similarly, Tv and Td are calculated only in the tumor compartment.

The aforementioned 11 compartments were divided into two sub-compartments: the vascular and extravascular spaces. Beyond these compartments, an extra compartment for blood circulation was added, and it did not divide into sub-compartments. For blood circulation and each sub-compartment, population conservation equations were formulated to capture the dynamics of every cell type. These equations describe the balance of cell populations, number of cells per volume, in terms of their accumulation in the volume of the compartment or sub-compartment, expressed as: cell accumulation = incoming fluxes - outgoing fluxes + transport across sub-compartments (vascular to extravascular space and vice versa) + other process such as cellular proliferation, death, cytolysis, phagocytosis, activation against tumor antigen, and deactivation of anti-tumor immune cells induced by regulatory cells (Tregs and M2).

The equations for the transport of dendritic cells through the vascular and extravascular space are presented below, and similar equations are also used for the other cells, with some extra terms which are presented later in Table 1.

The free circulating dendritic cells ($DC_{v_{j_F}}$, cells per volume) in the vascular space of the j-th compartment are given as per the equation (1) below:

$$V_{v_j} * \frac{dDC_{v_{j_F}}}{dt} = \sum_{\substack{k=organs \\ k\neq j,\ lnode}} \left( Q_k^{in} DC_{v_{k_F}}^{\ in} - Q_k^{out} DC_{v_{j_F}} \right) - J_{DC_j} * \left(1 - \frac{Im_{i_j}}{IM_{i_{max\,j}}}\right) * DC_{v_{j_A}} * V_{v_j} \qquad (1)$$

where the first term on the left side is the accumulation of $DC_{v_{j_F}}$ in the volume of the vascular space ($V_{v_j}$), the sum on the right side is the incoming minus the outgoing fluxes, where $Q_k^{in}$ is the blood flow from the k-th compartment and $Q_k^{out}$ is the outgoing blood flow from the j-th to the k-th compartment, which is multiplied by $DC_{v_{k_F}}^{in}$ (number of DC per volume in the k-th compartment) and $DC_{v_{j_F}}$ (number of DC per volume in the j-th compartment) respectively. The second term is the transmigration of DC from the vascular to the extravascular space, which depends on the transmigration constant ($J_{DC_j}$) and the level of immune cell saturation in the extravascular space of the j-th compartment, where $Im_{i_j}$ is the amount of immune cells and $IM_{i_{max_j}}$ is the maximum immune cells in the extravascular space of the j-th compartment.

The dendritic cells in the extravascular space ($DC_{i_j}$, cells per volume) of the j-th compartment are given as per the equation (2) below:

$$V_{i_j} * \frac{dDC_{i_j}}{dt} = J_{DC_j} * \left(1 - \frac{Im_{i_j}}{Im_{i_{max_j}}}\right) * DC_{v_{j_A}} * V_{v_j} - L_j * delta_{DC_j} * \left(1 - \frac{Im_{i_{lnode}}}{Im_{i_{max_{lnode}}}}\right) * DC_{i_j} + phi_{DC_j} * DC_{i_j} * \left(1 - \frac{Im_{i_j}}{Im_{i_{max_j}}}\right) * V_{i_j} \quad (2)$$

where the first term on the left side is the accumulation of $DC_{i_j}$ in the volume of the extravascular space ($V_{i_j}$), on the right side, the first term is the transmigration of dendritic cells from the vascular to extravascular space, the second term is migration to lymph nodes via lymphatic vessels, and the last term is the proliferation of $DC_{i_j}$. The transmigration term is the same as the equation (1) But with a positive sign in the equation (2). The migration to lymph nodes depends on the fluid absorption rate by the lymphatic vessels ($L_j$), the fraction ($delta_{DC_j}$) of $DC_{i_j}$ that can be recirculated, and the level of immune cell saturation in the lymph nodes. The proliferation term depends on the proliferation rate constant $phi_{DC_j}$ and the level of immune cell saturation in the extravascular space.

The dendritic cells in the extravascular space ($DC_{i_{lnode}}$, cells per volume) of the lymph node compartment are given as per the equation (3) below:

$$V_{i_{lnode}} * \frac{dDC_{i_{lnode}}}{dt} = J_{DC_{lnode}} * \left(1 - \frac{Im_{i_{lnode}}}{Im_{i_{max_{lnode}}}}\right) * DC_{v_{lnode_A}} * V_{v_{lnode}} + \sum_{k=organs} \left[L_k * delta_{DC_k} \left(1 - \frac{Im_{i_{lnode}}}{Im_{i_{max_{lnode}}}}\right) * DC_{i_k}\right] + phi_{DC_{lnode}} * DC_{i_{lnode}} * \left(1 - \frac{Im_{i_{lnode}}}{Im_{i_{max_{lnode}}}}\right) * V_{i_{lnode}} \quad (3)$$

where the first term on the left side is the accumulation of $DC_{i_{lnode}}$ in the volume of the extravascular space ($V_{i_{lnode}}$), on the right side, the first term is the transmigration of dendritic cells from the vascular to the extravascular space, the second term is the recirculation of $DC_{i_k}$ from the k-th compartments to the lymph node, and the last term is the proliferation of $DC_{i_{lnode}}$. The first and last terms on the right side of the equation (3) are similar to the equation (2) but the second term differs to consider the recirculation of dendritic cells from all compartments back to the lymph nodes.

The free dendritic cells ($DC_{v_{blood_F}}$, cells per volume) in the blood circulation are given as per the equation (4) below:

$$V_{v_{blood}} * \frac{dDC_{v_{blood_F}}}{dt} = \sum_{k=organs} \left(Q_k^{in} DC_{v_{k_F}}^{in} - Q_k^{out} DC_{v_{j_F}}\right) \quad (4)$$

where the first term on the left side is the accumulation of $DC_{v_{blood_F}}$ in the volume of the blood circulation ($V_{v_{blood}}$), and the sum on the right side is the incoming minus the outgoing fluxes, where $Q_k^{in}$ is the blood flow from the k-th compartment and $Q_k^{out}$ is the outgoing blood flow from the blood compartment to the k-th compartment, which is multiplied by $DC_{v_{k_F}}^{in}$ (number of DC per volume in the k-th compartment) and $DC_{v_{j_F}}$ (number of DC per volume in the j-th compartment) respectively.

In the PBPK model (equations 1-4), additional terms were incorporated into the population conservation equations for specific immune cell types and compartments to account for specific biological interactions. These terms reflect critical processes such as cell-to-cell interactions within the tumor microenvironment and other compartments. The terms that differentiate equations 1-4 for the different types of immune cells are presented in Table 1.

**Supplementary Table 1: Extra terms for the equations 1-4 for specific immune cell types and compartments to account for specific biological interactions.**

| Number of the equation where the term is added | Immune cell type for which the equation changed | j-th compartment where the term has effect | Extra term for the equations 1-6 for this immune cell and compartment |
|---|---|---|---|
| (1) | $APC_{v_{j_F}}$ | Bone, heart, kidney, liver, lung, muscle, skin, spleen, gastrointestinal tract, lymph nodes, tumor | $-(k_{sup_{M2_{APC}}} * M2_{v_{j_F}} + k_{sup_{Treg_{APC}}} * Treg_{v_{j_F}}) * APC_{v_{j_F}} * V_{v_j}$ |
| (2) | $APC_{i_j}$ | Bone, heart, kidney, liver, lung, muscle, skin, spleen, gastrointestinal tract, tumor | $-(k_{sup_{M2_{APC}}} * M2_{i_j} + k_{sup_{Treg_{APC}}} * Treg_{i_j}) * APC_{i_j} * V_{i_j}$ |
| (2) | $APC_{i_j}$ | Tumor | $+ x_{DC} * A_{c_{DC}} * \left(C_{i_{tumor}} + C_{i_{tumor_{dead}}}\right) * DC_{i_j} * V_{i_j} + x_{M1} * A_{c_{M1}} * (C_{i_{tumor}} + C_{i_{tumor_{dead}}}) * M1_{i_j} * V_{i_j}$ |

| | | | |
|---|---|---|---|
| (2) | $DC_{i_j}$ | Tumor | $-x_{DC} * A_{c_{DC}} * (C_{i_{tumor}} + C_{i_{tumor_{dead}}}) * DC_{i_j} * V_{i_{tumor}}$ |
| (2) | $M1_{i_j}$ | Tumor | $-x_{M1} * A_{c_{M1}} * (C_{i_{tumor}} + C_{i_{tumor_{dead}}}) * M1_{i_j} * V_{i_j} - x_{M1_{to_{M2}}} * C_{i_{tumor}} * M1_{i_j} * V_{i_j}$ |
| (2) | $M2_{i_j}$ | Tumor | $+x_{M1_{to_{M2}}} * C_{i_{tumor}} * M1_{i_j} * V_{i_j}$ |
| (1) | $TE1_{v_{j_F}}$ | Bone, heart, kidney, liver, lung, muscle, skin, spleen, gastrointestinal tract, lymph nodes, tumor | $-(k_{sup_{M2_{TE1}}} * M2_{v_{j_F}} + k_{sup_{Treg_{TE1}}} * Treg_{v_{j_F}}) * TE1_{v_{j_F}} * V_{v_j}$ |
| (2) | $TE1_{i_j}$ | Bone, heart, kidney, liver, lung, muscle, skin, spleen, gastrointestinal tract, tumor | $-(k_{sup_{M2_{TE1}}} * M2_{i_j} + k_{sup_{Treg_{TE1}}} * Treg_{i_j}) * TE1_{i_j} * V_{i_j}$ |
| (2) | $TE1_{i_j}$ | Lymph nodes | $+k_{TN} * TN_{i_j} * APC_{i_j} * V_{i_j}$ |
| (1) | $TE2_{v_{j_F}}$ | Bone, heart, kidney, liver, lung, muscle, skin, spleen, gastrointestinal tract, lymph nodes, tumor | $-(k_{sup_{M2_{TE2}}} * M2_{v_{j_F}} + k_{sup_{Treg_{TE2}}} * Treg_{v_{j_F}}) * TE2_{v_{j_F}} * V_{v_j}$ |
| (2) | $TE2_{i_j}$ | Bone, heart, kidney, liver, lung, muscle, skin, spleen, gastrointestinal tract, tumor | $-(k_{sup_{M2_{TE2}}} * M2_{i_j} + k_{sup_{Treg_{TE2}}} * Treg_{i_j}) * TE2_{i_j} * V_{i_j}$ |

Table 1 includes the suppression of APCs, TE1, and TE2 by M2 Macrophages and Tregs. For antigen-presenting cells (APCs), CD8+ effector T cells (TE1), and CAR-T cells (TE2), suppression terms were introduced across multiple organs in both extravascular spaces and vascular spaces (see Table 1). These terms account for the suppressive effects of M2 macrophages and regulatory T cells (Tregs) on the respective immune cell populations (e.g. $-(k_{sup_{M2_{APC\ or\ TE1}}} * M2_{i_j\ or\ v_j} + k_{sup_{Treg_{APC\ or\ TE1}}} * Treg_{i_j\ or\ v_j}) * Cell_{i_j\ or\ v_j} * V_{i_j\ or\ v_j}$ ). These terms reduce the populations of APCs, TE1, and TE2 in compartments where M2 macrophages and Tregs exert suppression.

Another term in Table 1 is related to tumor-specific activation of APC. Within the tumor compartment, an additional activation term was added for APCs in the extravascular space of the tumor compartment. This term accounts for the activation of APCs from dendritic cells (DCs) and M1 macrophages in response to tumor-associated antigens after phagocytosing both viable and dead tumor cells (e.g. $+x_{M1} * A_{c_{DC\ or\ M1}} * (C_{i_{tumor}} + C_{i_{tumor_{dead}}}) * Cell_{i_j} * V_{i_j}$). This term reflects the activation of APCs at the tumor site in response to immune-stimulatory signals and is added to the equation of APC and subtracted from the equation of DC or M1. Furthermore, M1 macrophages can transition into M2 macrophages under tumor-induced polarization with the term $x_{M1_{to\,M2}} * C_{i_{tumor}} * M1_{i_j} * V_{i_j}$ which is added to the equation of M2 and subtracted from the equation of M1.

The last extra term is related to the activation of CD8+ effector T cells (TE1) in lymph nodes. In the lymph node compartment, an additional term was added to the extravascular TE1 equation to reflect the activation of naive T cells ($TN_{i_j}$) by APCs, $k_{TN} * TN_{i_j} * APC_{i_j} * V_{i_j}$.

These extra terms enrich the model by introducing cell-specific interactions and biologically relevant processes, but also include the following 12 unknown parameters: suppression rate constant of APCs by M2 Macrophages $k_{sup_{M2_{APC}}}$, suppression rate constant of APCs by Treg $k_{sup_{Treg_{APC}}}$, suppression rate constant of CD8$^+$ effector T cells (TE1) and CAR-T cells (TE2) by M2 Macrophages $k_{sup_{M2_{TE1}}}$ and $k_{sup_{M2_{TE2}}}$, suppression rate constant of CD8$^+$ effector T cells (TE1) and CAR-T cells (TE2) by Tregs $k_{sup_{Treg_{TE1}}}$ and $k_{sup_{Treg_{TE2}}}$, activation ratio of DC to APC $x_{DC}$, phagocytosing rate constant of DC $A_{c_{DC}}$, activation ratio of M1 to APC $x_{M1}$, phagocytosing rate constant of M1 $A_{c_{M1}}$, transitioning rate constant of M1 to M2 induced by the tumor cells $x_{M1_{to\,M2}}$, activation rate constant of Naïve T cells to tumor antigen-specific effector T cells $k_{TN}$.

### Population conservation of cells in the tumor compartment

The population conservation equations of cells in the tumor compartment have the same formulation as described previously in equations 1-6, with the only difference on the left side of the equation (the accumulation) where the total volume and the density or concentration (number of cells per volume) of the cells changed simultaneously. For this reason, the accumulation was calculated for all cells based on their total number in the tumor compartment ($\frac{d\,(DC_{i_{tumor}} * V_{i_{tumor}})}{dt}$ or $\frac{d\,(DC_{v_{tumor}} * V_{v_{tumor}})}{dt}$ where $DC_{i_{tumor}} * V_{i_{tumor}}$ is the total number of DC in the tumor $n^{DC}_{i_{tumor}}$) and the density or concentration (cells per volume) of each cell was calculated by dividing the total number with the current tumor volume ($\frac{n^{DC}_{i_{tumor}}}{V_{i_{tumor}}}$).

The viable and dead tumor cells were calculated only in the extravascular space in the tumor compartment. The number of viable tumor cells in the extravascular space ($n^{c}_{i_{tumor}}$, total number of viable tumor cells) of the tumor compartment are given as per the equation (5) below:

$$\frac{dn^{c}_{i_{tumor}}}{dt} = \left( lg_{tumor} * C_{i_{tumor}} * \left(1 - \frac{Cell_{i_{tumor}}}{C_{tumor_{max}}}\right) - A_{c_{DC}} * C_{i_{tumor}} * \left(DC_{i_{tumor}} + APC_{i_{tumor}}\right) - A_{c_{M1}} * C_{i_{tumor}} * M1_{i_{tumor}} - C_{i_{tumor}} * \left(k_{rc1} * TE1_{i_{tumor}} + k_{rc2} * TE2_{i_{tumor}}\right) \right) * V_{i_{tumor}} \quad (5)$$

where the first term is the tumor cell proliferation which is proportional to the growth rate constant $lg_{tumor}$ and the density of viable cancer cells $C_{i_{tumor}}$ (cells per volume) and restricted with the levels of cells in extravascular space of the tumor, the second term is the phagocytosis of cancer cells by dendritic cells (DCs) and antigen-presenting cells (APCs), and the following term is phagocytosis of cancer cells by the macrophages (M1), and the last term is the cytolysis of cancer cells by cytotoxic effects of CD8+ effector T cells and CAR-T cells (TE1 and TE2).

The number of dead tumor cells in the extravascular space ($n^D_{i_{tumor}}$, total number of dead tumor cells) of the tumor compartment are given as per the equation (6) below:

$$\frac{dn^d_{i_{tumor}}}{dt} = \Big(-A_{c_{DC}} * D_{i_{tumor}} * \left(DC_{i_{tumor}} + APC_{i_{tumor}}\right) - A_{c_{M1}} * D_{i_{tumor}} * M1_{i_{tumor}} + C_{i_{tumor}} * \left(k_{rc1} * TE1_{i_{tumor}} + k_{rc2} * TE2_{i_{tumor}}\right)\Big) * V_{i_{tumor}} \quad (6)$$

where the first and second terms are the phagocytosis of dead cancer cells ($D_{i_{tumor}}$, cells per volume) by DC, APC, and M1 macrophages, and the last term is tumor cell death induced by effector CD8 T cells and CAR-T cells (TE1, TE2).

The tumor volume of the extravascular space $V_{i_{tumor}}$ was calculated based on the change in the total number of all cells in the extravascular space and the current density of the cells $Cell_{i_{tumor}}$ (total number of cells per volume) as per the equation (7) below.

$$\frac{dV_{i_{tumor}}}{dt} = \frac{\sum_{q=c,d,DC,APC,TE1,TE2,M1,M2,Treg} \frac{dn^q_{i_{tumor}}}{dt}}{Cell_{i_{tumor}}} \quad (7)$$

where the sum of the q-th cell type calculates the change in the total number of all cells in the tumor and the deviation by the current density of the cells ($Cell_{i_{tumor}}$) results in the change of volume ($\frac{dV_{i_{tumor}}}{dt}$).

The equations 7-9 introduce two extra unknown parameters, the growth rate constant $lg_{tumor}$, and the maximum density of cells in tumor compartment $C_{tumor_{max}}$.

### Estimation of Model Parameters

The optimization algorithm estimates the following parameters: tumor growth rate constant $lg_{tumor}$ and the initial tumor volume $V_{i_{tumor}0}$ which are related to the tumor cell line, other parameters related to the immune response such as the activation rate constant of naïve CD8$^+$ T cells $k_{TN}$, the recirculation rate of CD8$^+$ effector T cells $delta_{TE1_{lnode}}$, phagocytosing rate constant of cancer cells by DC, APC $A_{c_{DC}}$, and M1 $A_{c_{M1}}$, cytolytic rate constant of cancer cells by CD8$^+$ effector T cells $k_{rc1}$, and CAR-T cells $k_{rc2}$, transitioning rate constant of M1 to M2 induced by the tumor cells $x_{M1_{toM2}}$, suppression rate constant of CD8$^+$ effector T cells and CAR-T cells by M2 $k_{sup_{M2_{TE1}}}$ and $k_{sup_{M2_{TE2}}}$ and by Treg $k_{sup_{Treg_{TE1}}}$ and $k_{sup_{Treg_{TE2}}}$, suppression rate constant of APCs by M2 $k_{sup_{M2_{APC}}}$ and Treg $k_{sup_{Treg_{APC}}}$, proliferation rate constant of CAR-T cells under antigen stimulation in tumor $\varphi_{TE2_{tumor}}$, maximum time of CAR-T cells proliferation under antigen stimulation $t_{prol_{max}}$, and parameters related to the transmigration of immune cells to the tumor such as transmigration of Treg $J_{Treg_{tumor}}$, DC $J_{DC_{tumor}}$, M1 $J_{M1_{tumor}}$, M2 $J_{M2_{tumor}}$, CD8$^+$ effector T cells $J_{TE1_{tumor}}$, and CAR-T cells $J_{TE2_{tumor}}$.

For the murine GBM model, 15 parameters remained consistent across treatment groups ($lg_{tumor}$, $V_{i_{tumor}0}$, $delta_{TE1_{lnode}}$, $A_{c_{DC}}$, $A_{c_{M1}}$, $x_{M1_{to_{M2}}}$, $k_{sup_{M2_{TE1}}}$, $k_{sup_{Treg_{TE1}}}$, $k_{sup_{M2_{TE2}}}$, $k_{sup_{Treg_{TE2}}}$, $k_{sup_{M2_{APC}}}$, $k_{sup_{Treg_{APC}}}$, $\varphi_{TE2_{tumor}}$, $t_{prol_{max}}$, $k_{TN}$) while 8 parameters ($J_{Treg_{tumor}}$, $J_{DC_{tumor}}$, $J_{M1_{tumor}}$, $J_{M2_{tumor}}$, $J_{TE1_{tumor}}$, $J_{TE2_{tumor}}$, $k_{rc1}$, and $k_{rc2}$) varied between anti-VEGF and control treatments. In the human GBM model, the experiment was performed with immunodeficient mice and model parameters related to the adaptive immune system are not needed, thus 7 parameters ($lg_{tumor}$, $V_{i_{tumor}0}$, $A_{c_{DC}}$, $A_{c_{M1}}$, $x_{M1_{to_{M2}}}$, $k_{sup_{M2_{APC}}}$ and $k_{sup_{Treg_{APC}}}$) were consistent and 4 ($J_{Treg_{tumor}}$, $J_{DC_{tumor}}$, $J_{M1_{tumor}}$, and $J_{M2_{tumor}}$) differed between anti-VEGF and control treatments. The key parameters that were assumed to be affected by the anti-VEGF are the parameters which control the transmigration of immune cells in the tumor due to the improved normalized vasculature and the killing potential of T cells due to the immunosupportive tumor microenvironment caused by the vascular normalization. The resulting parameter estimates enabled the simulation of various therapeutic scenarios, ultimately providing insights into optimal treatment protocols for combining CAR-T cells therapy with anti-VEGF treatment for GBM. The Supplementary Table 2 presents the model parameters that were calculated by the optimization process and are not affected by anti-VEGF normalization; and Supplementary Table 3 presents the parameters that are affected by anti-VEGF.

**Supplementary Table 2: Parameters estimated by optimization that are not affected by anti-VEGF (constant across treatments)**

| Parameter | Description | Value for GSC005 | Value for MGG8 |
|---|---|---|---|
| $lg_{tumor}$ | Tumor growth rate constant | 1.05 [1/d] | 0.754 [1/d] |
| $V_{i_{tumor0}}$ | Initial tumor volume | 0.00222 [$mm^3$] | 9.1264e-05 [$mm^3$] |
| $delta_{TE1_{lnode}}$ | Recirculation rate of CD8$^+$ effector T cells | 0.00119 [1] | - |
| $A_{c_{DC}}$ | Phagocytosis rate constant by dendritic cells/APCs | 0.27016 [$cm^3$/d] | 0.48747 [$cm^3$/d] |
| $A_{c_{M1}}$ | Phagocytosis rate constant by M1 macrophages | 95.7797 [$cm^3$/d] | 0.081193 [$cm^3$/d] |
| $x_{M1_{to_{M2}}}$ | Transition rate of M1 to M2 macrophages | 7.51e-06 [$cm^3$/d] | 2.5384e-06 [$cm^3$/d] |
| $k_{sup_{M2_{TE1}}}$ | Suppression of CD8+ effector T cells by M2 macrophages | 3.78 [$cm^3$/d] | - |
| $k_{sup_{M2_{TE2}}}$ | Suppression of CAR-T cells by M2 macrophages | 0.0981 [$cm^3$/d] | - |
| $k_{sup_{Treg_{TE1}}}$ | Suppression of CD8+ effector T cells by Tregs | 4.35e-08 [$cm^3$/d] | - |
| $k_{sup_{Treg_{TE2}}}$ | Suppression of CAR-T cells by Tregs | 2.33e-06 [$cm^3$/d] | - |
| $k_{sup_{M2_{APC}}}$ | Suppression of APCs by M2 macrophages | 2.65 [$cm^3$/d] | 2.65 [$cm^3$/d] |
| $k_{sup_{Treg_{APC}}}$ | Suppression of APCs by Tregs | 1.86 [$cm^3$/d] | 1.86 [$cm^3$/d] |
| $\varphi_{TE2_{tumor}}$ | CAR-T proliferation rate under antigen stimulation | 4.59e-09 [$cm^3$/d] | - |

| | | | |
|---|---|---|---|
| $t_{prol_{max}}$ | Maximum duration of CAR-T proliferation under antigen stimulation | 1.59[d] | - |
| $k_{TN}$ | Activation rate constant of naïve $CD8^+$ T cells | 2970 [$cm^3$/d] | - |

**Supplementary Table 3: Parameters estimated by optimization that are affected by anti-VEGF (differ across treatments with and without anti-VEGF)**

| Parameter | Description | Value without anti-VEGF | | Value with anti-VEGF | |
|---|---|---|---|---|---|
| | | GSC005 | MGG8 | GSC005 | MGG8 |
| $J_{Treg_{tumor}}$ | Transmigration of Tregs into tumor | 6024.8322 [1/d] | 1011.9995 [1/d] | 1991.6031 [1/d] | 6.3721 [1/d] |
| $J_{DC_{tumor}}$ | Transmigration of dendritic cells into tumor | 1.6617e-05 [1/d] | 0.072353 [1/d] | 0.00065133 [1/d] | 0.47446 [1/d] |
| $J_{M1_{tumor}}$ | Transmigration of M1 macrophages into tumor | 1.01 [1/d] | 1.0562 [1/d] | 1.59 [1/d] | 1.0929 [1/d] |
| $J_{M2_{tumor}}$ | Transmigration of M2 macrophages into tumor | 1.0578e-06 [1/d] | 0.00025436 [1/d] | 4.44e-07 [1/d] | 7.451e-10 [1/d] |
| $J_{TE1_{tumor}}$ | Transmigration of $CD8^+$ effector T cells into tumor | 14.671 [1/d] | - | 69.0017 [1/d] | - |
| $J_{TE2_{tumor}}$ | Transmigration of CAR-T cells into tumor | 0.65546 [1/d] | - | 1.8168 [1/d] | - |
| $k_{rc1}$ | Cytolytic rate of tumor cells by effector $CD8^+$ T cells | 1.5302e-05 [$cm^3$/d] | - | 3.5601e-06 [$cm^3$/d] | - |
| $k_{rc2}$ | Cytolytic rate of tumor cells by CAR-T cells | 1.3227e-06 [$cm^3$/d] | - | 1.6623e-06 [$cm^3$/d] | - |

### PBPK model parameters taken from literature

The rest parameters of the PBPK model are physiological and literature-derived and are presented in Supplementary Table 4. Supplementary Table 4 presents only the parameters that were not zero and their values were derived from [1,2].

**Supplementary Table 4: Physiological and literature-derived model parameters.**

| Category | Parameter | Value | Description |
|---|---|---|---|
| **Blood flows** | $Q_{blood}$ | 4.38 [$cm^3$/min] | Blood flow rate |
| | $Q_{bone}$ | 0.17 [$cm^3$/min] | Bone blood flow |
| | $Q_{heart}$ | 0.28 [$cm^3$/min] | Heart blood flow |
| | $Q_{kidney}$ | 0.80 [$cm^3$/min] | Kidney blood flow |
| | $Q_{liver}$ | 1.10 [$cm^3$/min] | Liver blood flow |
| | $Q_{liver_{in}}$ | Q_liver+Q_gi-L_gi+Q_spleen-L_spleen | Effective liver inflow |
| | $Q_{muscle}$ | 0.80 [$cm^3$/min] | Muscle blood flow |
| | $Q_{skin}$ | 1.21 [$cm^3$/min] | Skin blood flow |

| | | | |
|---|---|---|---|
| | $Q_{spleen}$ | 0.05 [cm³/min] | Spleen blood flow |
| | $Q_{gi}$ | 0.90 [cm³/min] | GI tract blood flow |
| | $Q_{lnode}$ | 0.05 [cm³/min] | Lymph node blood flow |
| **Lymph flows** | $L_{blood}$ | 0 [cm³/min] | Blood lymph flow |
| | $L_{bone}$ | $6\times10^{-5}$ [cm³/min] | Bone lymph flow |
| | $L_{heart}$ | $1\times10^{-5}$ [cm³/min] | Heart lymph flow |
| | $L_{kidney}$ | $1.7\times10^{-4}$ [cm³/min] | Kidney lymph flow |
| | $L_{liver}$ | $2\times10^{-4}$ [cm³/min] | Liver lymph flow |
| | $L_{lung}$ | $1\times10^{-4}$ [cm³/min] | Lung lymph flow |
| | $L_{muscle}$ | $6\times10^{-4}$ [cm³/min] | Muscle lymph flow |
| | $L_{skin}$ | $1\times10^{-5}$ [cm³/min] | Skin lymph flow |
| | $L_{spleen}$ | $2\times10^{-6}$ [cm³/min] | Spleen lymph flow |
| | $L_{gi}$ | $7\times10^{-4}$ [cm³/min] | GI lymph flow |
| | $L_{lnode}$ | $1.8\times10^{-3}$ [cm³/min] | Lymph node lymph flow |
| **Vascular volumes** | $V_{v_{blood}}$ | 0.774 [cm³] | Blood vascular volume |
| | $V_{v_{bone}}$ | 0.080 [cm³] | Bone vascular volume |
| | $V_{v_{heart}}$ | 0.007 [cm³] | Heart vascular volume |
| | $V_{v_{kidney}}$ | 0.030 [cm³] | Kidney vascular volume |
| | $V_{v_{liver}}$ | 0.095 [cm³] | Liver vascular volume |
| | $V_{v_{lung}}$ | 0.019 [cm³] | Lung vascular volume |
| | $V_{v_{muscle}}$ | 0.150 [cm³] | Muscle vascular volume |
| | $V_{v_{skin}}$ | 0.200 [cm³] | Skin vascular volume |
| | $V_{v_{spleen}}$ | 0.010 [cm³] | Spleen vascular volume |
| | $V_{v_{gi}}$ | 0.100 [cm³] | GI vascular volume |
| | $V_{v_{lnode}}$ | 0.010 [cm³] | Lymph node vascular volume |
| **Interstitial volumes** | $V_{i_{blood}}$ | 0 [cm³] | Blood interstitial volume |
| | $V_{i_{bone}}$ | 0.280 [cm³] | Bone interstitial volume |
| | $V_{i_{heart}}$ | 0.019 [cm³] | Heart interstitial volume |
| | $V_{i_{kidney}}$ | 0.101 [cm³] | Kidney interstitial volume |
| | $V_{i_{liver}}$ | 0.190 [cm³] | Liver interstitial volume |
| | $V_{i_{lung}}$ | 0.057 [cm³] | Lung interstitial volume |
| | $V_{i_{muscle}}$ | 1.032 [cm³] | Muscle interstitial volume |
| | $V_{i_{skin}}$ | 0.999 [cm³] | Skin interstitial volume |
| | $V_{i_{spleen}}$ | 0.020 [cm³] | Spleen interstitial volume |

| | | | |
|---|---|---|---|
| | $V_{i_{gi}}$ | 0.600 [cm³] | GI interstitial volume |
| | $V_{i_{lnode}}$ | 0.020 [cm³] | Lymph node interstitial volume |
| **Immune cell trafficking – APCs** | $J_{APC_{liver}}$ | $2.9\times10^{-3}$ [1/min] | APC trafficking to liver |
| | $J_{APC_{lung}}$ | $2.9\times10^{-3}$ [1/min] | APC trafficking to lung |
| | $J_{APC_{spleen}}$ | $2.9\times10^{-3}$ [1/min] | APC trafficking to spleen |
| | $J_{APC_{lnode}}$ | $2.9\times10^{-3}$ [1/min] | APC trafficking to lymph node |
| | $J_{APC_{tumor}}$ | $2.9\times10^{-3}$ [1/min] | APC trafficking to tumor |
| | $delta_{APC_{liver}}$ | 1 | Fraction of APCs in liver |
| | $delta_{APC_{lung}}$ | 1 | Fraction of APCs in lung |
| | $delta_{APC_{spleen}}$ | 1 | Fraction of APCs in spleen |
| | $delta_{APC_{lnode}}$ | $1.8\times10^{-8}$ | Fraction of APCs in lymph node |
| | $delta_{APC_{tumor}}$ | 0.1 | Fraction of APCs in tumor |
| **Immune cell trafficking – DCs** | $J_{DC_{liver}}$ | $2.9\times10^{-3}$ [1/min] | DC trafficking to liver |
| | $J_{DC_{lung}}$ | $2.9\times10^{-3}$ [1/min] | DC trafficking to lung |
| | $J_{DC_{spleen}}$ | $2.9\times10^{-3}$ [1/min] | DC trafficking to spleen |
| | $delta_{DC_{bone}}$ | 1 | Fraction of DCs in bone |
| | $delta_{DC_{liver}}$ | 1 | Fraction of DCs in liver |
| | $delta_{DC_{lung}}$ | 1 | Fraction of DCs in lung |
| | $delta_{DC_{spleen}}$ | 1 | Fraction of DCs in spleen |
| | $delta_{DC_{lnode}}$ | 1 | Fraction of DCs in lymph node |
| | $delta_{DC_{tumor}}$ | 0.1 | Fraction of DCs in tumor |
| | $phi_{DC_{bone}}$ | 6.2 [1/d] | DC proliferation rate in bone |
| | $x_{DC}$ | 0.75 | DC differentiation fraction |
| **Immune cell trafficking – M1 macrophages** | $J_{M1_{liver}}$ | $2.9\times10^{-3}$ [1/min] | M1 trafficking to liver |
| | $J_{M1_{lung}}$ | $2.9\times10^{-3}$ [1/min] | M1 trafficking to lung |
| | $J_{M1_{spleen}}$ | $2.9\times10^{-3}$ [1/min] | M1 trafficking to spleen |
| | $delta_{M1_{bone}}$ | 1 | Fraction of M1 in bone |

| | | | |
|---|---|---|---|
| | $delta_{M1_{liver}}$ | 1 | Fraction of M1 in liver |
| | $delta_{M1_{lung}}$ | 1 | Fraction of M1 in lung |
| | $delta_{M1_{spleen}}$ | 1 | Fraction of M1 in spleen |
| | $delta_{M1_{lnode}}$ | 1 | Fraction of M1 in lymph node |
| | $delta_{M1_{tumor}}$ | 0.1 | Fraction of M1 in tumor |
| | $phi_{M1_{bone}}$ | 6.2 [1/d] | M1 proliferation in bone |
| **Immune cell trafficking – M2 macrophages** | $J_{M2_{liver}}$ | 2.9×10⁻³ [1/min] | M2 trafficking to liver |
| | $J_{M2_{lung}}$ | 2.9×10⁻³ [1/min] | M2 trafficking to lung |
| | $J_{M2_{spleen}}$ | 2.9×10⁻³ [1/min] | M2 trafficking to spleen |
| | $x_{M1}$ | 0.75 | M1→M2 differentiation fraction |
| | $delta_{M2_{bone}}$ | 1 | Fraction of M2 in bone |
| | $delta_{M2_{liver}}$ | 1 | Fraction of M2 in liver |
| | $delta_{M2_{lung}}$ | 1 | Fraction of M2 in lung |
| | $delta_{M2_{spleen}}$ | 1 | Fraction of M2 in spleen |
| | $delta_{M2_{lnode}}$ | 1 | Fraction of M2 in lymph node |
| | $delta_{M2_{tumor}}$ | 0.1 | Fraction of M2 in tumor |
| | $phi_{M2_{bone}}$ | 0.86 [1/d] | M2 proliferation in bone |
| **Immune cell trafficking – Tregs** | $J_{Treg_{liver}}$ | 2.9×10⁻³ [1/min] | Treg trafficking to liver |
| | $J_{Treg_{lung}}$ | 2.9×10⁻³ [1/min] | Treg trafficking to lung |
| | $J_{Treg_{spleen}}$ | 2.9×10⁻³ [1/min] | Treg trafficking to spleen |
| | $delta_{Treg_{bone}}$ | 1 | Fraction of Tregs in bone |
| | $delta_{Treg_{liver}}$ | 1 | Fraction of Tregs in liver |
| | $delta_{Treg_{lung}}$ | 1 | Fraction of Tregs in lung |
| | $delta_{Treg_{spleen}}$ | 1 | Fraction of Tregs in spleen |
| | $delta_{Treg_{lnode}}$ | 1 | Fraction of Tregs in lymph node |
| | $phi_{Treg_{blood}}$ | 1.736×10⁻² [cm³/d] | Treg proliferation rate |

| | | | |
|---|---|---|---|
| **Immune cell trafficking – CD8$^+$ T cells (TE1)** | $J_{TE1_{liver}}$ | 2.9×10$^{-3}$ [1/min] | Effector CD8$^+$ T trafficking to liver |
| | $J_{TE1_{lung}}$ | 2.9×10$^{-3}$ [1/min] | Effector CD8$^+$ T trafficking to lung |
| | $J_{TE1_{spleen}}$ | 2.9×10$^{-3}$ [1/min] | Effector CD8$^+$ T trafficking to spleen |
| | $delta_{TE1_{liver}}$ | 1 | Fraction of effector T in liver |
| | $delta_{TE1_{lung}}$ | 1 | Fraction of effector T in lung |
| | $delta_{TE1_{spleen}}$ | 1 | Fraction of effector T in spleen |
| **Immune cell trafficking – CAR-T (TE2)** | $J_{TE2_{liver}}$ | 2.9×10$^{-3}$ [1/min] | CAR-T trafficking to liver |
| | $J_{TE2_{lung}}$ | 2.9×10$^{-3}$ [1/min] | CAR-T trafficking to lung |
| | $J_{TE2_{spleen}}$ | 2.9×10$^{-3}$ [1/min] | CAR-T trafficking to spleen |
| | $delta_{TE2_{liver}}$ | 1 | Fraction of CAR-T in liver |
| | $delta_{TE2_{lung}}$ | 1 | Fraction of CAR-T in lung |
| | $delta_{TE2_{spleen}}$ | 1 | Fraction of CAR-T in spleen |

### Model validation with other relevant data from the literature [3,4]

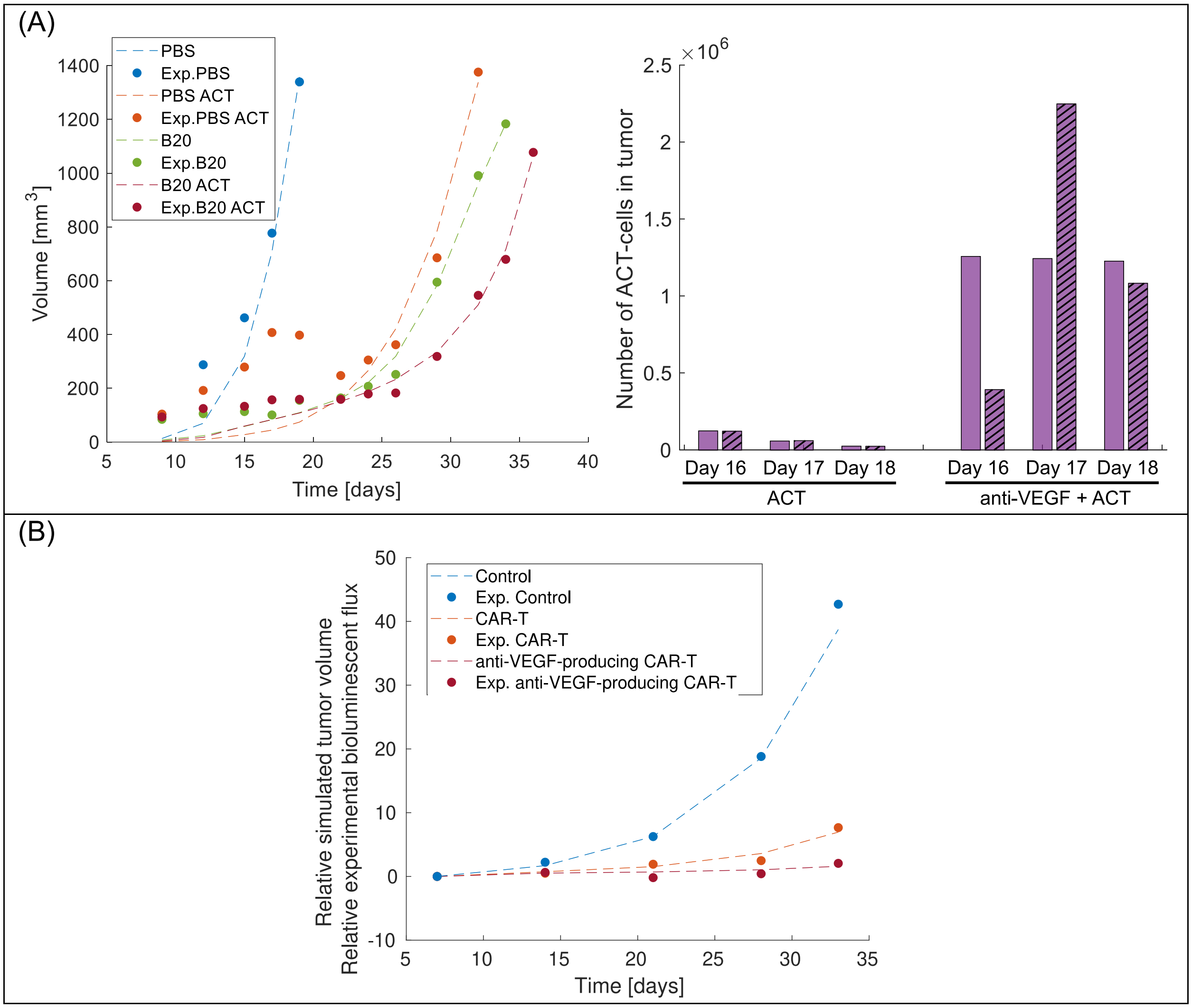


**Supplementary Figure 1: Model validation with other relevant data. A) Model validation with adoptive T cell therapy in melanoma murine tumors from [4]. B) Model validation with anti-VEGF producing CAR-T cells therapy in ovarian murine tumors from [3].**
